# Copepodamides in marine and freshwater copepods-*similar but different*

**DOI:** 10.1101/2023.11.13.566526

**Authors:** Sina Arnoldt, Milad Pourdanandeh, Ingvar Spikkeland, Mats X. Andersson, Erik Selander

## Abstract

Marine copepods, the most abundant group of zooplankton in the worlds oceans, imprint their surrounding waters with chemical cues, called copepodamides. Copepodamides induce defensive traits such as toxin production, bioluminescence, and colony size plasticity in a variety of marine phytoplankton. The role of copepodamides in freshwater ecosystems is, however, unknown. Here we report the consistent presence of copepodamides in copepods from six Swedish lakes. Copepodamide concentrations in freshwater copepods are similar to those of marine copepods, around 0.1 ppt of dry mass in millimetre sized individuals. The composition substantially overlaps with marine copepodamides but is also distinctly different. Freshwater copepods are largely dominated by one of the two subgroups of copepodamides, the dihydro-copepodamides, whereas marine copepods more commonly contain representatives from both subgroups. We describe 10 new copepodamide structures, four of which only found in freshwater copepods. Taxonomic groups had consistent copepodamide profiles across sampling sites and timepoints, supporting the presence of species specific copepodamide signatures. The presence of copepodamides in freshwater environments warrants study into their potential function as predator alarm cues also in freshwater systems.

## Introduction

Phytoplankton sense chemical alarm cues from zooplankton predators and respond by inducing a range of defensive traits. These include morphological, behavioural, chemical, and life history responses [1-4]. Two groups of cueing compounds have been identified so far. Aliphatic sulphates or sulfamates from freshwater cladocerans, a group of short chain (9-11 carbon) aliphatic compounds, sometimes branched, with different level of saturation that induce colony formation [5] in the freshwater green algae *Desmodesmus subspicatus* (previously *Scenedesmus subspicatus*). Marine copepods produce the other known class of defence inducing compounds, copepodamides, a group of structurally closely related taurine conjugated lipids (Figure 1). Copepodamides, hereafter referring to the general group of compounds, are divided into two subgroups based on the presence of a methyl (dihydro-copepodamides, hereafter referred to as “dhCA”) or methylene group (copepodamides, hereinafter “CA”) in position C3. The fatty acid attached to C5 is variable. Long chain polyunsaturated ω-3 fatty acid such as docosahexaenoic (22:6), eicosapentaenoic (20:5), or stearidonic (18:4) are common in marine copepodamides [6]. Other fatty acids, however, including even-numbered saturated fatty acids such as myristic (14:0) or palmitic (16:0) acid are also found. 31 copepodamide derivatives have been described thus far [7]. Copepodamide concentrations are correlated to copepod densities in the ocean and reach bioactive levels when copepods are abundant [8]. Harmful algal bloom (HAB) forming phycotoxin producers such as *Pseudo-nitzschia* spp. and *Alexandrium* spp. respond by producing more amnesic [9] and paralytic shellfish toxins. Bioluminescent taxa such as *Lingulodinium polyedrum* and *Alexandrium tamarense* increase light production [10]. Chain forming diatoms split up colonies into smaller units [11,12]. Other diatoms have been shown to increase in silica content and stickiness, which results in cell aggregation [13]. Induced toxin production and morphological changes are accompanied by increased resistance to grazers [10,12,14,15]. Well defended taxa may subsequently benefit from a competitive edge which can contribute to the formation of HABs.

**Figure 1.**
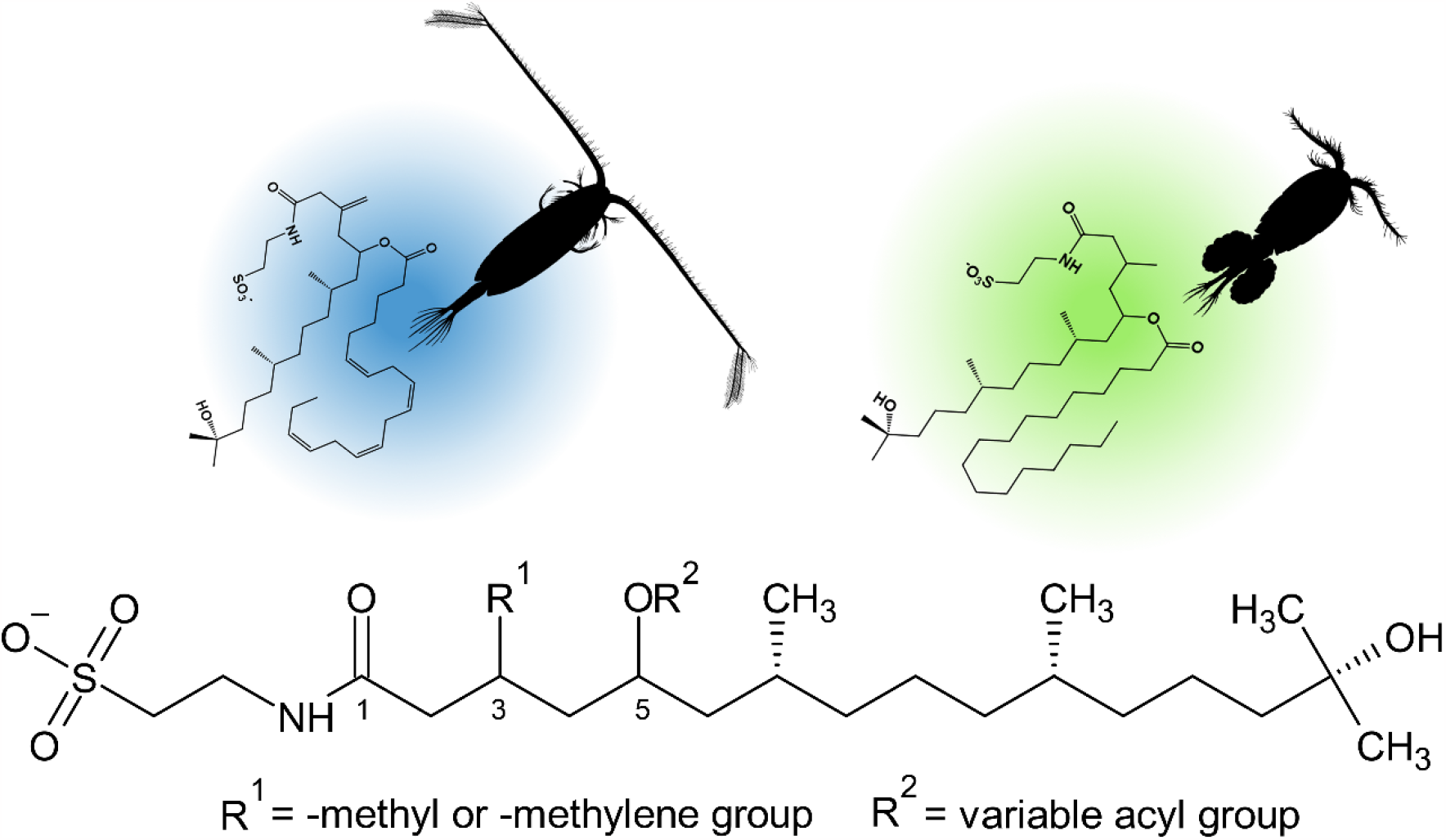
General structure of copepodamides. Two main subgroups exist, determined by the presence of methylene (copepodamide/CA, blue) or methyl (dihydro-copepodamide/dhCA) at R^1^. The blend is species specific, but the fatty acid side chain (at position R^2^) changes with diet. Copepodamides are named by the acyl group [7] followed by the scaffold name e.g. 22:6 dihydro-copepodamide for a dhCA scaffold with a docosahexaenoic acyl group in position R^2^. Silhouettes are of a calanoid (left) and cyclopoid (right) copepod.

Incorporating copepod densities, or direct measurements of copepodamides in mussels, can indeed improve the lead time and precision in HAB forecasting models [16]. Copepodamides are universally present in planktic marine cyclopoid and calanoid copepods. Given the dominance of copepod zooplankton in all oceans, copepodamides are likely among the most widespread chemical alarm cues known.

Aliphatic sulphates have only been reported from limnic environments, whereas copepodamides are, except from a single measurement in a pond in Gothenburg, Sweden [7], only reported from the ocean. There are, however, empirical evidence that limnic dinoflagellates also respond to chemical cues from copepod grazers. Resting stages of freshwater dinoflagellates of the genera *Ceratium* and *Peridinium* delay excystment when copepods are abundant in the water column above [17]. The identity of the cueing compound(s) is not known, and the role of copepodamides in limnic ecosystems remains to be explored.

Here we prove the presence of copepodamides in freshwater copepods from five lakes located on the Swedish west coast and one northern latitude ice-covered lake. We analyse bulk zooplankton samples and extracts from individual copepods to compare the composition and amounts of freshwater copepodamides with marine samples.

## Materials and methods

### Collection of copepods

Six lakes/ponds and four marine sites were sampled (Figure 2, for more information about the sampling sites see Table S.1). Freshwater copepods were collected with a handheld plankton net (mesh size: 65 μm, diameter: 15 cm), except for the ice-covered lake (F6) which was sampled through the ice with a smaller net (mesh size: 25 µm, diameter: 10 cm). Marine copepods were sampled with the same handheld plankton net at sampling sites M2 and M4, with a WP2 net (mesh size: 90 µm, diameter 57 cm diameter) onboard R/V Oscar von Sydow at M1, and with a handheld net (mesh size 200 µm, diameter: 57 cm) from a small outboard motorised boat at M3. Horizontal/oblique net tows at 1-2 m depth (average) were carried out at all sampling stations, except for M1, M3, and F6. Vertical tows were used at M1 and M3, from 25 m and 50 m depth to the surface respectively. Vertical/oblique tows were also used to sample through the ∼60 cm diameter drilled ice-hole at lake F6. Individual copepods were sorted under a stereomicroscope (Stemi-305 Zeiss) within 0.5 to 9 h after sampling (median = 2.5h). Copepods were transferred to a 1.5 mL HPLC glass vial and water carefully removed with a pasteur pipette. The remaining copepod was extracted in 1 mL methanol overnight at - 20 °C. The sample was concentrated by evaporating the methanol under a stream of nitrogen (40 °C) and resolved in 50 μL methanol and stored at −20 °C until analysis. Copepods were identified to the lowest possible taxonomic level and prosome length measured under an inverted microscope (Axiovert A1, Zeiss). Biomass (dry weight) was estimated from collated length-weight regressions (Table. S2).

**Figure 2.**
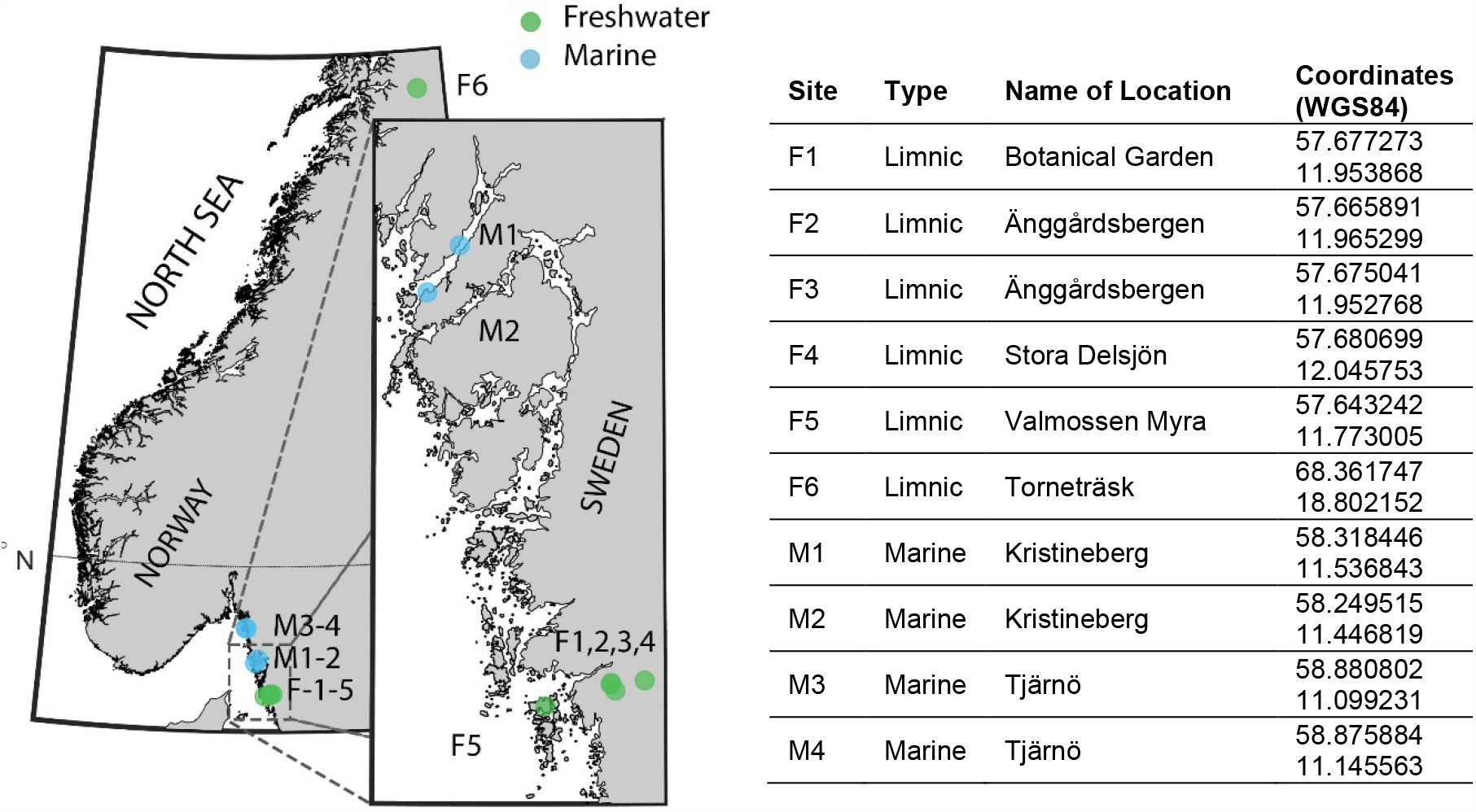
Map of sampling stations with names and positions. Freshwater sites are shown in green and marine in blue. Stations F1-F3 markers overlap, F5 is located on the island Brännö. M3 & M4 markers overlap and are in the Koster Fjord.

### Copepodamide analysis

Copepodamides were analysed on an Agilent 1260 HPLC system coupled to an Agilent 6475 triple quadrupole MS detector. Copepodamides were separated on a Prevail C18, 3 μm, 2.1*150 mm column thermostated to 50 °C using a gradient elution from 95% eluent A (methanol:acetonitrile:water 35:35:30), 5 % isopropanol (eluent B) to 83 % B over 18 min followed by a 7 min re-equilibration 0.2 % formic acid and 0.1 % ammonia was added to both eluents. Bulk zooplankton samples containing hundreds of copepods were scanned for compounds of 600 to 1000 m/z producing the product ions characteristic of CAs (430.3 m/z) or dhCAs (432.3 m/z, Supplementary Data. 1). Copepodamides in single copepod extracts were analysed with multiple reaction monitoring (MRM) experiments using transitions making up ≥80% of the total ion counts from the precursor scans in addition to the ten known copepodamides to maintain sensitivity (Supplementary Data. 2). Copepodamide measurements were calibrated against authentic copepodamide standards assuming the same ionisation efficiency for all copepodamides. For details on this, chromatography, and source settings, see Selander and colleagues [6].

### Multivariate ordinations and statistical analyses

Copepodamide composition of marine and freshwater samples were compared using permutational multivariate analysis of variance (PERMANOVA) [18] in the R package *vegan* [19]. Homogeneity of variance was tested using a permutation test of multivariate dispersion (PERMDISP) [20]. Non-metric multidimensional scaling (nMDS) was used to visualise the multivariate data in two-dimensional space. Difference in copepodamide content between freshwater and marine copepods was assessed with an analysis of covariance (ANCOVA), controlling for their body size (estimated dry mass). Assumptions of (i) linearity were assessed visually and with linear regression, (ii) homogeneity of regression slopes by evaluating the covariate-predictor interaction, and (iii) conditional normality was tested with Shapiro-Wilks’ test visual assessment. The data was non-normally distributed, but ANCOVA (and ANOVAs in general) procedures are robust against violations of normality when sample sizes are large and similar among groups [21]. Bray-Curtis distance [22] was used as dissimilarity metric for all nMDS, PERMANOVA and PERMDISP procedures, permutations were set to n = 9999 and significance level was set to a = 0.05 for all tests. All statistical analyses were performed in R version 4.2.3 [23] via RStudio [24]. Supplementary figures S1-S3 and tables S1-S3 are available in Supplementary information.

### Code availability and supplementary information

The R code, and the output it generated, used for this article is available as a HTML-file (Supplementary Code). This, and the source R-markdown file, is also available in the public repository linked to this article [25], see section “Data Availability” below for details.

## Results

The freshwater samples were dominated by cyclopoids (Figure. S1); *Cyclops strenuus* was found in almost all samples and contributed 47% of the individually extracted freshwater copepods. Other unidentified cyclopoids were the second largest group (23 %) and *Macrocyclops albidus* found in a single lake. Freshwater calanoids were only sampled and extracted from the ice-covered lake F6 where *Eudiaptomus graciloides* was found. Marine samples, in contrast, were dominated by calanoids (Figure. S2); *Temora longicornis* was the most abundant of the individually extracted copepods (34%), followed by *Centropages hamatus* (16%) and *Calanus* sp. (16%).

In total we detected 35 putative copepodamides, 11 were present in both marine and freshwater samples, 6 only in freshwater, and 18 only in marine samples (Figure 3A-B, Table 1). Fragmentation experiments of abundant compounds revealed the presence of fragments matching dhCA, (C_22_H_42_NO_5_S m/z 432.28) and CA scaffolds (C_22_H_40_NO_5_S m/z 430.26), taurine (C_2_H_6_NO_3_S m/z 124.01), sulfonate (SO_3_ m/z 79.96) as well as similar retention times to that of known copepodamides. Freshwater copepods almost exclusively contained dhCAs. Trace amounts of CA were found in individual copepods from the lake on the island Brännö (F5) and Stora Delsjön (F4, Figure 4). Marine samples contained comparably more CAs (Figure 3B). Table 1 contains a complete list of copepodamides with the putative identity of the acyl group inferred from the neutral loss in MS-MS experiments. Copepodamides from freshwater samples appear to be more variable in saturation level, see for example the sequence of 18:5-, 18:4-, 18:3-, and 18:2-dhCA (Figure 3A, Table 1). Moreover, limnic copepodamides include more odd number fatty acids such as C_15_ and C_17_. Copepodamides in marine samples, on the other hand, were dominated by polyunsaturated ω-3 fatty acids. Ten of the putative copepodamide structures found have not previously been described (Table 1).

**Table 1.** List of putative copepodamides in freshwater and marine bulk zooplankton samples with fatty acid identity inferred from the neutral loss associated with loss of the acyl group. Presence in marine (M), freshwater (F) or both (B) is indicated. Asterisk (*) indicate compounds found in only one sample.

| Scaffold | Putative fatty acyl | $m/z$ precursor ion | Described in | Present in |
| --- | --- | --- | --- | --- |
| Dihydro-copepodamides<br>($m/z$ product ion: 432) | 14:0 | 660.5 | Grebner 2019 | B |
|  | 15:1 | 672.5 | This study | F |
|  | 15:0 | 674.5 | This study | B |
|  | 16:4 | 680.5 | This study | M |
|  | 16:3 | 682.5 | This study | M |
|  | 16:2 | 684.5 | This study | B |
|  | 16:1 | 686.5 | This study | B |
|  | 16:0 | 688.6 | Grebner 2019 | B |
|  | 17:1 | 700.5 | This study | F |
|  | 17:0 | 702.7 | This study | F |
|  | 18:5 | 706.5 | Grebner 2019 | M |
|  | 18:4 | 708.6 | Grebner 2019 | B |
|  | 18:3 | 710.6 | Grebner 2019 | B |
|  | 18:2 | 712.6 | Grebner 2019 | B |
|  | 18:1 | 714.6 | Grebner 2019 | B |
|  | 20:5 | 734.6 | Grebner 2019 | B |
|  | 20:4 | 736.6 | Grebner 2019 | F |
|  | 20:3 | 738.7 | This study | F |
|  | 20:1 | 742.6 | This study | M |
|  | 22:6 | 760.6 | Grebner 2019 | B |
|  | 22:5 | 762.6 | Grebner 2019 | F |
| Copepodamides<br>($m/z$ product ion: 430) | 14:0 | 658.5 | Grebner 2019 | M |
|  | 16:4* | 678.5 | Grebner 2019 | M |
|  | 16:1 | 684.4 | Grebner 2019 | M |
|  | 16:0 | 686.6 | Grebner 2019 | M |
|  | 18:5 | 704.5 | Grebner 2019 | M |
|  | 18:4 | 706.6 | Grebner 2019 | M |
|  | 18:3 | 708.6 | Grebner 2019 | M |
|  | 18:2 | 710.6 | Grebner 2019 | M |
|  | 18:1 | 712.6 | Grebner 2019 | M |
|  | 18:0 | 714.6 | Grebner 2019 | M |
|  | 20:5 | 732.6 | Grebner 2019 | M |
|  | 20:4 | 734.6 | Grebner 2019 | M |
|  | 22:6 | 758.7 | Grebner 2019 | M |
|  | 22:5 | 760.6 | Grebner 2019 | M |

**Figure 3.**
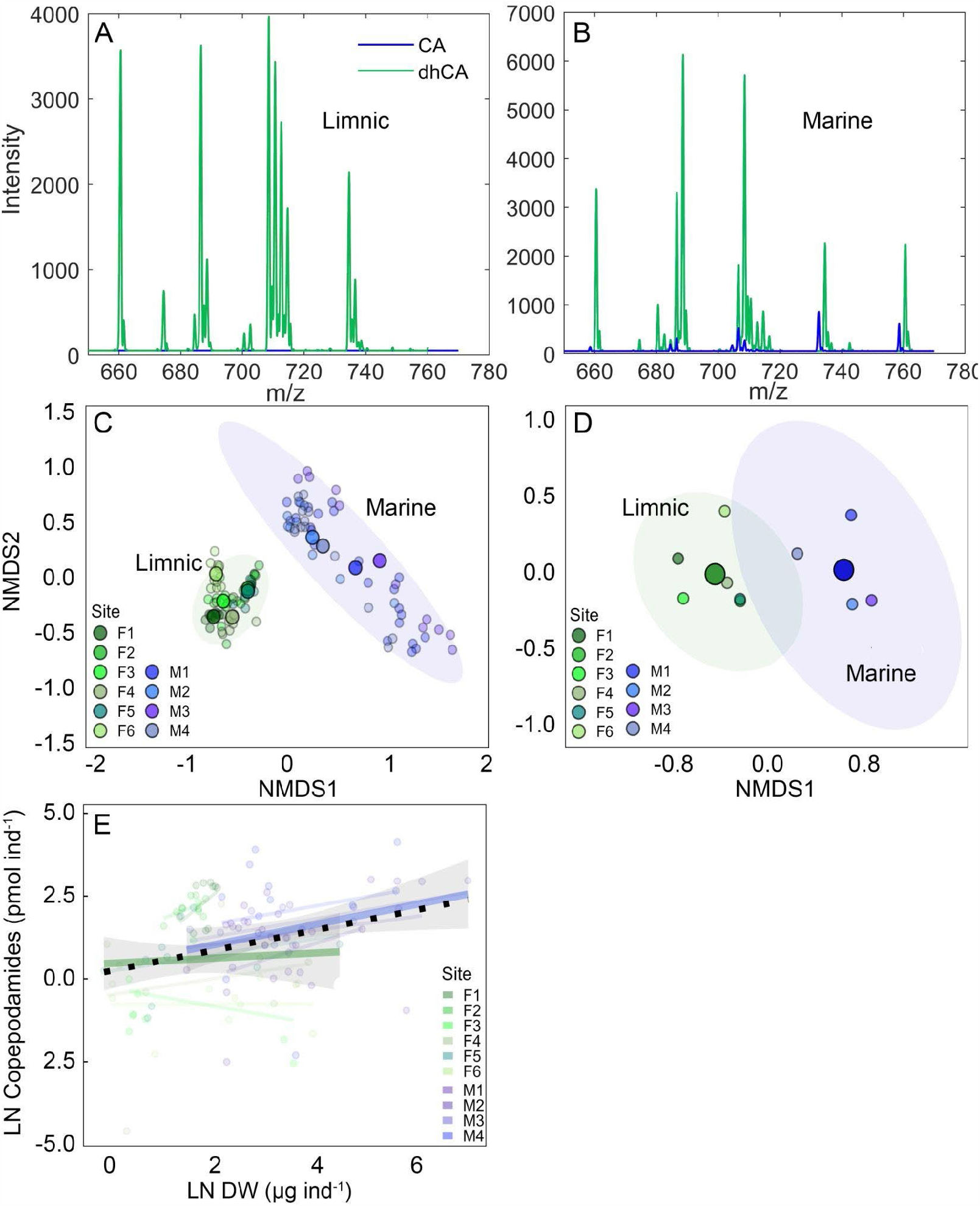
A: Representative mass spectra from precursor ion scans of copepodamides in bulk zooplankton samples from limnic (F5) and **B**: marine (M4). Note the absence of copepodamides (CA, blue) in the limnic samples. **C**: Non-metric dimensional scaling (nMDS) ordination of copepodamide composition in individual copepods (smaller points) and **D**: bulk zooplankton samples. Larger points denote the centroid for each group, coloured ellipses are 95% confidence intervals of centroids. Stress values for the nMDS models: 0.11 and 0.04 respectively. **E**: Total copepodamide content in individual copepods (pmol ind^-1^, filled circles) plotted against their estimated dry mass (µg). Thin lines denote regression lines for the individual sampling sites, thicker coloured lines denote regression lines for the two habitat types and dotted black line denotes the global regression (Ln(CA pmol) = 0.296 * Ln(dry mass µg) + 0.25, R^2^ = 0.087). Shaded error bands denote 95% confidence intervals for the habitat regressions.

**Figure 4.**
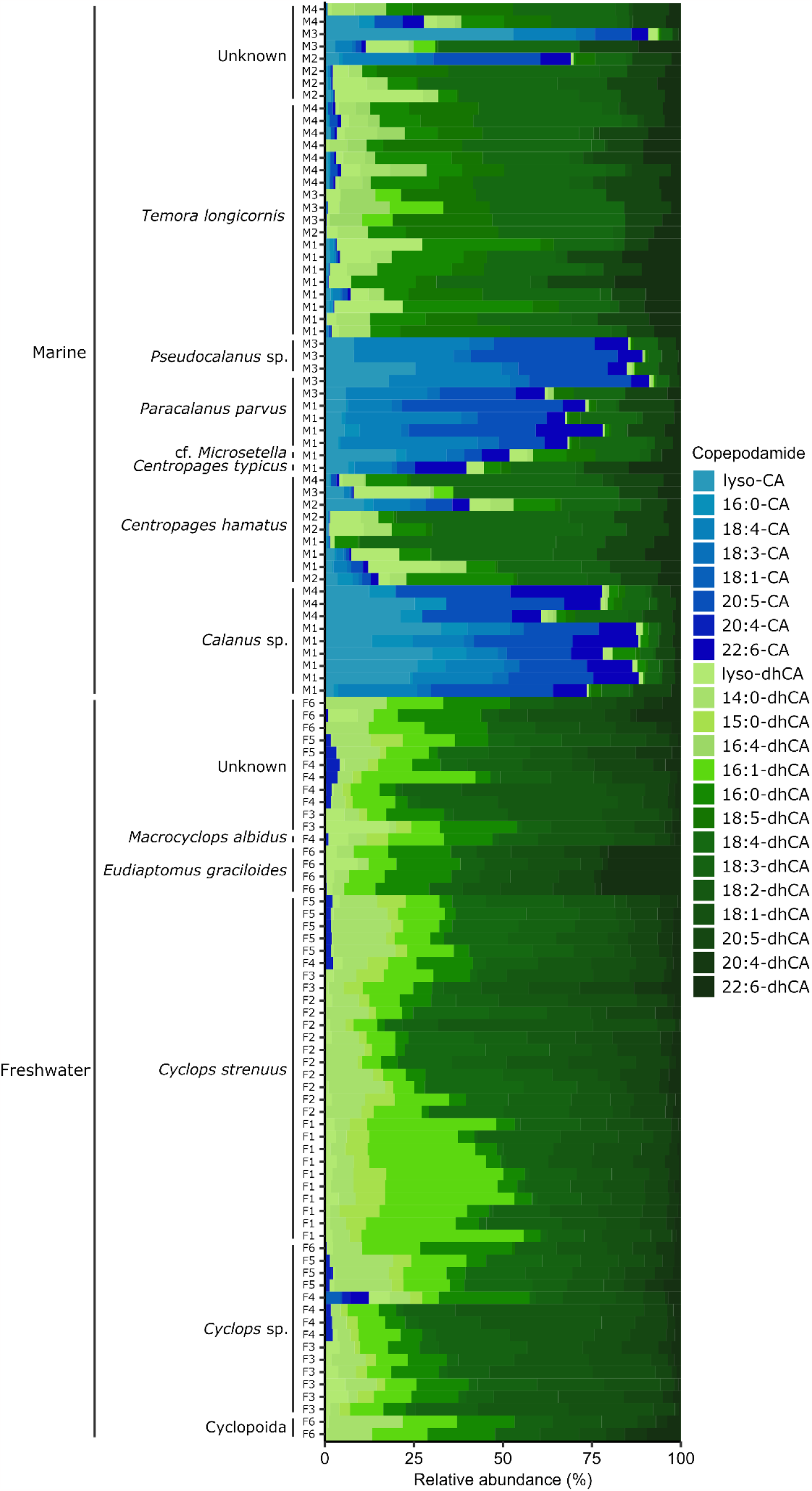
Composition of copepodamides in individual copepods from targeted LC-MS analysis of the most abundant copepodamides. Organised by habitat (marine & freshwater), taxon and sampling site. Blue and green bars denote CAs and dhCAs respectively.

The composition of copepodamides differed significantly between freshwater and marine copepods for both the targeted analysis of individually extracted copepods (PERMANOVA: df = 1, 102, p < 0.001, Figure 3C) and non-targeted analysis of bulk zooplankton samples (PERMANOVA: df = 1, 8, p = 0.004, Figure 3D). Copepods of the same taxonomic unit had similar copepodamide signatures across sampling stations and sampling times (Figure 4). Taxonomy explained 48% of the variance left unexplained by the other model terms (partial eta squared: η_p_^2^), and habitat (freshwater or marine) explained 18%. Copepodamide content in individually extracted copepods was comparable in similarly sized marine and freshwater copepods (ANCOVA: df = 1, 112, p = 0.14, η_p_^2^ = 0.019; Figure 3E). Larger copepods contained more copepodamides than smaller individuals (Linear regression: p = 0.003), but less in proportion to their mass. A millimetre sized copepod (100 µg dry weight) contained around 10 ng copepodamides, or 0.1 ppt of dry mass.

## Discussion

Copepodamides were present in all freshwater samples as well as in all individually extracted freshwater copepods, which shows that they are generally present also in freshwater copepods. The amount is comparable to that of similar sized marine copepods (Figure 3C). but varied an order of magnitude between samples from different lakes. Copepods from lakes F6 and F3 e.g., had 5 to 10 times lower levels of copepodamides than the lakes F1-2 and F4-5 (Figure 3). F6 was the only frozen and snow-covered lake we sampled from. Light penetration and consequently primary production is limited in these conditions, this food-limitation likely contributed to the low copepodamide concentrations in lake F6. Copepodamides are hypothesised to be involved in digestion [6], as the composition reflects the recent diet of the copepods [7]. Structurally related compounds have been suggested to be related to digestion in protozoans [26] and in several animals [27]. Different levels of gut fullness is therefore likely to affect copepodamide content in addition to composition. The reason for the low levels in lake F3 is less obvious. It is possible that the recent feeding history and nutritional state of the copepods contributed to this variation, just as copepodamide composition is known to change with diet. The half-life of copepodamides in seawater is short, around 3 h at 15 ºC in seawater [8] but longer, around 35 h when bound in mussel tissue [16]. Differences in the handling time from sampling to extraction may consequently contribute to the observed differences. However, Lakes F6 and F3 were processed after 2.4 and 3.3 h, which is longer than F1 and F2 (0.9 and 1.9 h) but not F5 (3.1 h) which was the lake with the highest copepodamide content per copepod. Moreover, the slight decrease in size-adjusted copepodamide content as a function of time was non-significant (Multiple regression: p = 0.062, Figure. S3 & Table. S3). This suggests that the variation in copepodamide content was likely due to differences in phylogeny, size, or sites, rather than to processing time.

Copepodamide composition in marine and freshwater copepods clearly overlap, as approximately one third of the copepodamides were found in both marine and freshwater copepods. Yet, the remaining two thirds were restricted to marine or limnic copepods. Copepodamides from freshwater copepods were dominated by dhCAs, with low levels of CAs detected only in lakes F4 and F5. Lake F5 is situated on an island and is possibly more affected by the ocean than the other lakes, e.g., in terms of salt deposition. Lake F4, in contrast, is 12 km from the sea and is to be considered fully limnic. dhCAs are an order of magnitude more potent toxin inducers than CAs in the dinoflagellate *Alexandrium minutum* [6], which suggest that higher proportion of dhCAs may be of ecological significance. The fatty acyl moiety in position C5 changes with diet within days in the marine copepod *Temora longicornis*, but the ratio of dhCAs to CAs appears more stable [7].

The saturation level of the fatty acid moiety appears more variable in limnic samples, and limnic samples also include more odd number fatty acids such as C_15_ and C_17_ commonly found in prokaryotes [28]. The inherent difference in fatty acid composition in marine and limnic system is probably the main reason for the discrepancy between marine and freshwater copepodamides [29]. C_18_ fatty acids with different degrees of saturation are common in limnic seston [30], and the relative abundance changes with decreasing salinity in e.g. the green algae *Chlorella vulgaris* and *Acutodesmus obliquus* [31]. The copepodamide composition is closely related to the taxonomic affinity and appears relatively stable across locations and time points (Figure 4). This is surprising considering the rapid dynamics of the phytoplankton community composition in time and space, which suggests that copepodamide profile is stabilised either by selective feeding or species-specific differences in lipid metabolism that preserve differences between taxa. Freshwater samples were dominated by cyclopoid copepods, whereas marine samples mainly consisted of calanoids. This phylogenetic bias may have contributed to the difference between marine and freshwater copepodamide compositions. However, the freshwater calanoid evaluated here (*Eudiaptomus graciloides*) clearly clustered with the freshwater cyclopoids rather than the marine calanoids, suggesting that the difference in copepodamide compositionis not primarily a consequence of phylogenetic bias. Analysis of copepodamide composition in more marine cyclopoids and limnic calanoids would help to shed light on this query.

We add the 10 new copepodamide structures to the 31 previously reported (Table 1). Only 4 of in total 41 copepodamides were only found in freshwater samples. The full flat structure is only known for 10 out of the 41, and the putative structures listed here require further interrogation to establish full structure and chirality. From an ecological point of view the partitioning between dhCAs and CAs is likely more important than differences in the fatty acid moiety which, as long as there is a fatty acid attached, seems to be of less importance for the structure-activity relationship [6,8]

In conclusion we find that copepodamides are common in freshwater copepods. There is a substantial overlap with marine copepods, yet limnic copepodamides are distinct in terms of the near complete dominance of dihydro-copepodamides and a different composition of the acyl groups. The generality of copepodamides as alarm signals in the ocean suggests that their possible role as kairomones also in freshwater environments should be studied.

## Acknowledgements

We thank Kristie Rigby, Isabel Casties, Lars Ljungqvist, and Linda Svanberg for their help with sampling marine copepods. This work was financially supported by a Swedish Research Councils grant to E.S (VR 2019-05238)

## Author contributions

E.S and M.P conceived the study. S.A, E.S, and E.S planned the field sampling, which S.A conducted. E.S, S.A, M.X.A analysed the copepodamides. E.S and I.S conducted taxonomic identification of copepods. M.P conducted formal analysis and visualisation. S.A, M.P, and E.S wrote the manuscript with input from all authors. All authors have read and approved the final manuscript.

## Data availability

All data and analysis code generated or used in this article is stored in a public Zenodo respiratory (DOI: 10.5281/zenodo.8047945) (Activated upon acceptance)

## Additional information

The authors declare no competing interests.

