## Supplementary information for "Copepodamides in marine and freshwater copepods-*similar but different*"

**for**

### *-similar but different*

Sina Arnoldt[1+], Milad Pourdanandeh[1+], Ingvar Spikkeland[2], Mats X. Andersson[3], Erik Selander[4*]

1: Department of Marine Sciences, University of Gothenburg, Medicinaregatan 7B, SE41390, Gothenburg, Sweden

2: Østfold Museum Foundation, Sarpsborg, Østfold, Norway

3: Department of Biological and Environmental Sciences, University of Gothenburg, Medicinaregatan 7B, SE41390, Gothenburg, Sweden

4: Department of Biology - Aquatic Ecology, Lund University, Sölvegatan 37, SE22362, Lund, Sweden

[+] Shared first authors

Public repository containing all source datasets, analysis datasets, code, and output: (DOI) 10.5281/zenodo.8047945

**Table S1.** Description of sampling sites.

| Site | Habitat | Name of Location | Coordinates (WGS84) | Type | Sampling | Description |
|---|---|---|---|---|---|---|
| F1 | Limnic | Botanical Garden, Gothenburg | 57.677273 11.953868 | Artificial pond | 65µm net, 15 cm diameter, 1-2 metre depth horizontal/oblique tow | A small artificial pond in the Gothenburg Botanical Gardens, approximately 30-40 $m^2$, depth of 1-2 m |
| F2 | Limnic | Trindemossen, Änggårdsbergen, Gothenburg | 57.665891 11.965299 | Natural lake | 65µm net, 15 cm diameter, 1-2 metre depth horizontal/oblique tow | A small lake in Änggårdsbergen, Gothenburg, approximately 7 200 $m^2$, unknown maximum and average depth, high productivity, clear water |
| F3 | Limnic | Finnsmossen, Änggårdsbergen, Gothenburg | 57.675041 11.952768 | Natural lake | 65µm net, 15 cm diameter, 1-2 metre depth horizontal/oblique tow | A small lake in Änggårdsbergen, Gothenburg, approximately 12 500 $m^2$, unknown maximum and average depth, high productivity, low visibility, |
| F4 | Limnic | Stora Delsjön, Gothenburg | 57.680699 12.045753 | Natural lake | 65µm net, 15 cm diameter, 1-2 metre depth horizontal/oblique tow | One of the larger lakes in Gothenburg, popular for recreational activities. Area: 1.2 $km^2$. Depth: 5.2 m (average), 23 m (max). Good ecological status, high nutrient conditions, poor oxygenation in some parts, fair chemical status (except for levels of Hg and PBDEs). Source: WISS database, Swedish Water District Authorities and County Administrative Boards (https://viss.lansstyrelsen.se/) |
| F5 | Limnic | Valmossen, Brännö | 57.643242 11.773005 | Natural lake | 65µm net, 15 cm diameter, 1-2 metre depth horizontal/oblique tow | Small natural lake on the island Brännö outside Gothenburg. Approx. 7 400 $m^2$, unknown depth, high productivity |
| F6 | Limnic | Torneträsk, Abisko | 68.361747 18.802152 | Natural lake | 25µm net, 15 cm diameter, 1-2 metre depth horizontal/oblique tow | One of Sweden's largest lakes, located in Kiruna municipality in the north of Sweden. Area: 330 $km^2$, Depth: 51.8 m (average), 168 m (max), good ecological status, high nutrient levels, fair chemical status (except for levels of Hg and PBDEs). Sources: Swedish Meteorological and Hydrological Institute (https://www.smhi.se/data/hydrologi/svenskt-vattenarkiv) & WISS database, Swedish Water District Authorities and County Administrative Boards (https://viss.lansstyrelsen.se/) |
| M1 | Marine | Gullmarn Fjord, Fiskebäckskil | 58.318446 11.536843 | Fjord | 90µm net, 57 cm diameter, vertical tow from 25 m depth to surface, onboard R/V Oscar von Sydow | Gullmarn Fjord, between Torseröd and Alsbäck |
| M2 | Marine | Kristineberg, Fiskebäckskil | 58.249515 11.446819 | Near harbour | 65µm net, 15 cm diameter, 1-2 metre depth horizontal/oblique tow | Close to the harbour of Kristineberg Center for Marine Research and Innovation. |
| M3 | Marine | Koster Fjord, Strömstad | 58.880802 11.099231 | Fjord | 200 µm net, x cm diameter, vertical tow from ~50m to surface, onboard a small outboard motorised boat. | Deep water (~200 m) site between Saltö and the Koster Islands |
| M4 | Marine | Tjärnö, Strömstad | 58.875884 11.145563 | Shallow cove, near harbour | 65µm net, 15 cm diameter, 1-2 metre depth horizontal/oblique tow | Shallow (5-10 m), close to the harbour of Tjärnö Marine Laboratory. |

**Table. S2** Average estimates of terms used to estimate dry mass (g) from prosome length for different copepod taxa. Collated from published literature (below conversion equation) and averaged for the full range of prosome lengths measured in the primary sources for each taxon. Detailed calculations and regression plots are available in the public repository linked to this publication (File name: Masterfile_targeted_data_final.xlsx, Sheets: E-H).

| Taxa | a | b |
|---|---|---|
| Cyclopoida | 6.614E-06 | 2.465 |
| Calanoida | 2.801E-05 | 3.079 |
| Temora | 3.130E-05 | 3.060 |
| Micosetella | 2.239E-10 | 1.150 |
| Grand mean (used for unkown taxa) | 1.731E-04 | 2.798 |

Dry mass (g) = **a** x prosome length(mm)^**b**

| Taxa | Source |
|---|---|
| *Acartia clausi* | Hay 1991 |
| *Calanus finmarchicus* | Hay 1991 |
| *Calanus helgolandicus* | Hay 1991 |
| *Metridia lucens* | Hay 1991 |
| *Centropages typicus* | Hay 1991 |
| *Centropages hamatus* | Breteler et al. 1982 - Growth and devolpment of four calanoid copepod species under experimental and natural conditions |
| *Paracalanus parvus* | Hay 1991 |
| *Pseudocalanus elongatus* | Hay 1991 |
| *Acartia clausi* | Breteler et al. 1982 |
| *Temora longicornis* | Hay 1991 |
| *Aanthocyclops vernalis* | Rosen 1981 - Length-Dry Weight Relationships of Some Freshwater Zooplankton |
| Mesocyclops edax | Rosen 1981 |
| *Tropocyclops prasinus* | Castillo-Noll 2007 - Length-weight relationships for zooplanktonic species of a tropical Brazilian lake: Lake Monte Alegre. |
| Cyclopoida (inc. females with eggs) | Dumont et al 1975 - The Dry Weight Estimate of Biomass in a Selection of Cladocera, Copepoda and Rotifera from the Plankton Periphyton and Benthos of Continental Waters |
| Cyclopoida (excl. females with eggs) | Dumont et al 1975 |
| Pooled cyclopoida | Burgis (in press) in Dumont et al 1975 - The Dry Weight Estimate of Biomass in a Selection of Cladocera, Copepoda and Rotifera from the Plankton Periphyton and Benthos of Continental Waters |
| Cyclopoida 3 species C1-C6 | Nakamura et al 2017 - Length-weight Relationships and Chemical Composition of the Dominant Mesozooplankton Taxa/species in the Subarctic Pacific, with Special Reference to the Effect of Lipid Accumulation in Copepoda |
| *Tropocyclops prasinus* | Castillo-Noll 2007 |
| *Microsetella norvegica* | Satapoomin 1999 - Carbon content of some common tropical Andaman Sea copepods |

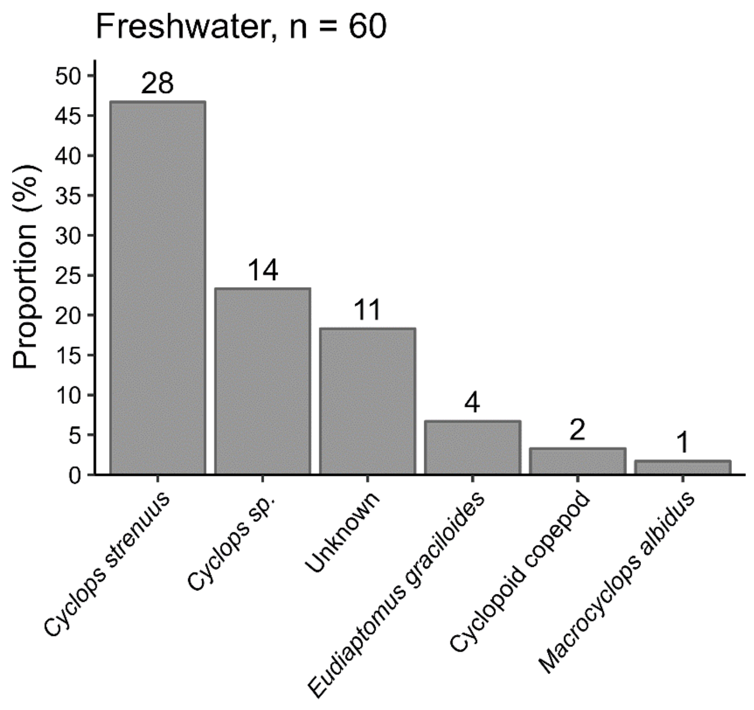


**Figure. S1** Taxonomic composition (%) of n = 60 individual copepods, sampled from six freshwater sites in Sweden. Values above bars are counts.

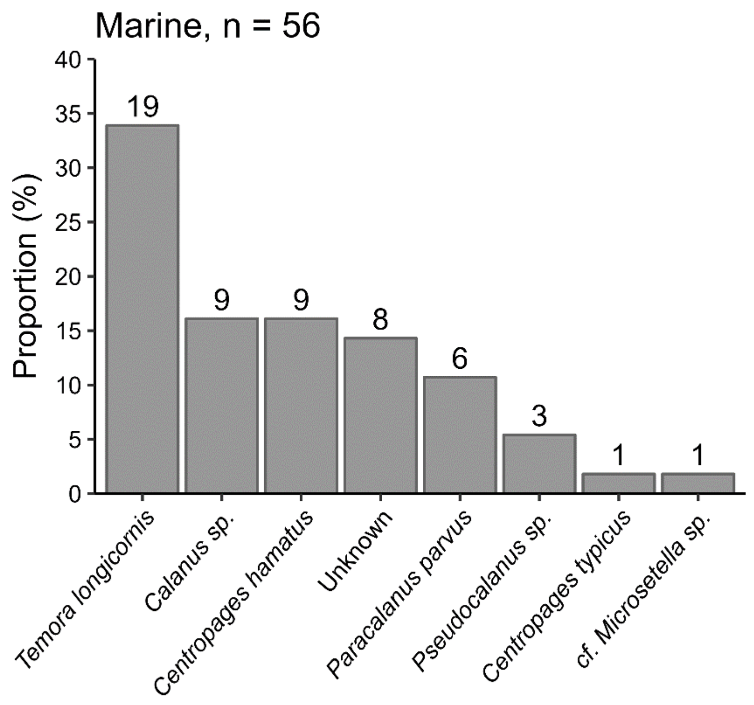


**Figure. S2** Taxonomic composition (%) of n = 56 individual copepods, sampled from four marine sites in Sweden. Values above bars are counts.

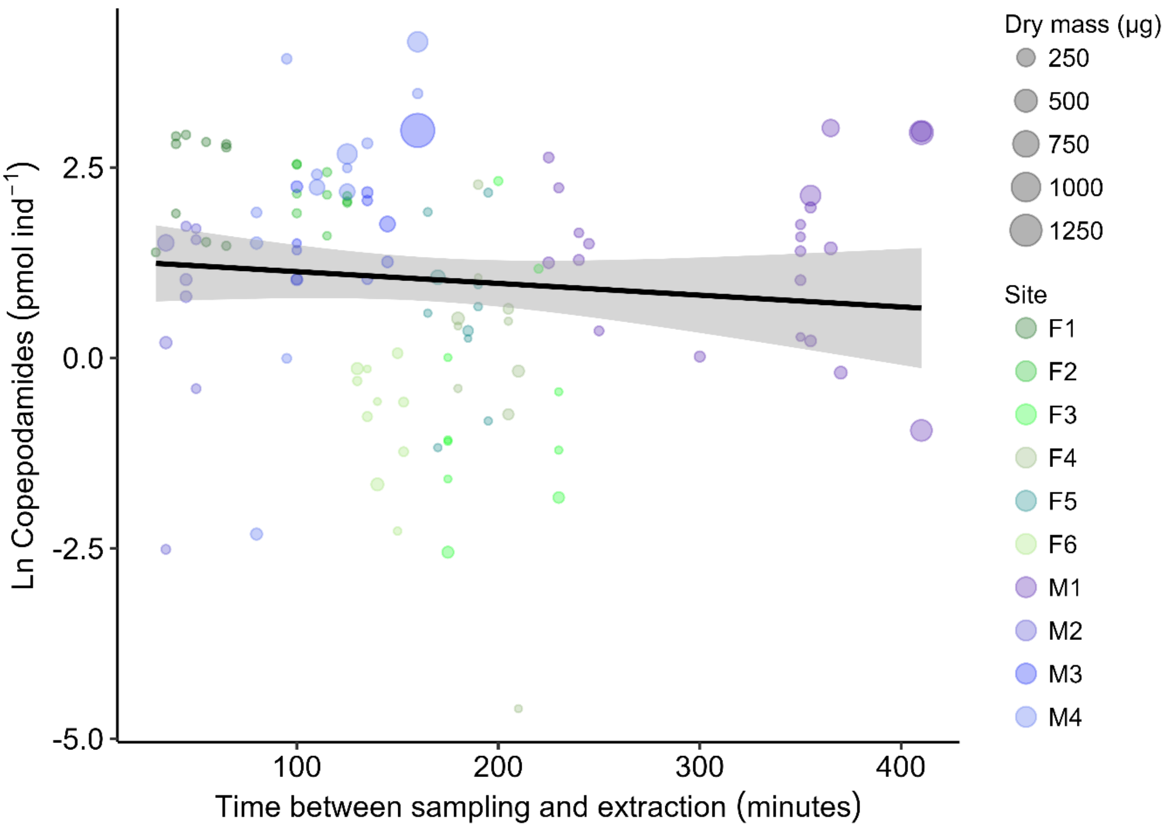


**Figure. S3** Relationship between copepodamide content (on a natural log scale) of individual copepods and the time between sampling and extraction. Points are coloured by sampling site (greens = freshwater , blues/purples = marine) and size scaled by the dry mass of the copepod. The black line is the least square regression line (y = -0.002x + 1.33, $R^2$ = 0.014) and the shaded band is the 95% confidence interval of the regression line.

**Table. S3** Linear regression models of copepodamide content in individual copepods (on a natural log scale) as a function of the time between sampling and extraction, adjusted for the dry mass of the copepods (left) and non-adjusted (right).

| | **Corrected for copepod mass** | | | **Raw time effect** | | |
|---|---|---|---|---|---|---|
| **Variable** | **Beta** | **95% CI**[1] | **p-value** | **Beta** | **95% CI**[1] | **p-value** |
| **Intercept** | 1.3378 | 0.7712, 1.9044 | **<0.001** | 1.3294 | 0.7459, 1.9130 | **<0.001** |
| **Time (min)** | -0.0029 | -0.0059, 0.0001 | 0.062 | -0.0020 | -0.0050, 0.0011 | 0.2 |
| **Dry mass (μg)** | 0.0026 | 0.0008, 0.0044 | **0.006** | | | |
| $R^2$ | 0.079 | | | 0.014 | | |
| df | 2 | | | 1 | | |
| Residual df | 113 | | | 114 | | |

[1] CI = Confidence Interval
