## Supplementary code for "Copepodamides in marine and freshwater copepods-*similar but different*": Supplementary Code.html

##### Author: Milad Pourdanandeh

#### 2023-11-06

#### Purpose:

This code generates statistical analyses, multivariate ordinations
and visualisations of copepodamides from marine and freshwater copepods,
used in the research article titled below.

#### Article:

“Copepodamides in marine and freshwater copepods - *similar but
different*”

##### DOI:

10.5281/zenodo.8047945

##### Article authors:

**Sina Arnoldt (co-first author)**, **Milad
Pourdanandeh (co-first author)**, Ingvar Spikkeland, Mats X.
Andersson & Erik Selander (*corresponding author*)

#### About the data:

Two data files are used in this analysis,
*Arnoldt\_targeted\_analysis\_data.csv* and
*Arnoldt\_untargeted\_analysis\_data.csv*, derived from the source
files *Masterfile\_targeted\_data\_final.xlsx* and
*Precursor\_data\_Deisotoped.xlsx* respectively. The analysis files
are also published as **Supplementary Data. 1** &
**Supplementary Data. 2** in the article. For details about
the two source files, see their respective READMEs (first sheet in each
file). All files are publicly available for download at Zenodo (DOI:
10.5281/zenodo.8047945).

#### Targeted data:

The targeted LC-MS input data
(*Arnoldt\_targeted\_analysis\_data.csv*) consists of columns:

- **Sample** - Unique identification for each
  sample.
- **Type** - Habitat type, marine or freshwater.
- **Site** - Sampling site, six freshwater sites (F1-F4)
  and four marine ones (M1-M4).
- **Time\_diff\_h** - Time between sampling and extraction
  in hours for each sample.
- **Time\_diff\_min** Time between sampling and extraction
  in minutes for each sample.
- **No\_of\_copepods** - Number of copepods in the sample,
  all should be 1 in all.
- **Taxon** - Taxonomic information for each copepod,
  freshwater copepods identified by Ingvar Spikkeland, marine copepods by
  Erik Selander.
- **Sex** - Sex of the copepods.
- **Stage** - The putative developmental stage of the
  copepods.
- **Prosome\_length\_µm** - Length of the copepods prosome
  in µm.
- **Dry\_weight\_g** - Estimated dry mass, derived from
  prosome length, in g.
- **Dry\_weight\_µg** - Estimated dry mass, derived from
  prosome length, in µg.
- **Dry\_weight\_ng** - Estimated dry mass, derived from
  prosome length, in ng.
- **Data\_file** - The mass spectrometer data file
  containing the data fro each sample.
- **Acq\_Date\_Time** - The date and time that data was
  aquired from the LC-MS
- **CA\_per\_copepod\_mol** - Estimate of the total amount
  of copepodamides for each sample (in mol).
- **CA\_per\_copepod\_pmol**Estimate of the total amount of
  copepodamides for each sample (in pmol).
- **448.3\_CA - 758.6\_CA** - Concentrations (in µM) of
  copepodamides (with 430 *m/z*scaffold).
- **450.3\_dhCA - 760.5\_dhCA** - Concentrations (in µM) of
  dihydro-copepodamides (with 432 *m/z* scaffold).

#### Non-targeted/precursor data:

The non-targeted LC-MS input data
(*Arnoldt\_untargeted\_analysis\_data.csv*) consists of columns:

- **Type** - Habitat type, marine or freshwater.
- **Site** - Sampling site, six freshwater sites (F1-F4)
  and four marine ones (M1-M4).
- **660.5\_dhCA - 762.6\_dhCA** - Each dihydro-copepodamide
  as a proportion of total coppeodamides.
- **658.5\_CA - 760.7\_CA** - Each copepodamide (with the
  430 *m/z* copepodamide scaffold) as a proportion of total
  copepodamides.

### Preparation

```
# Set the working directory to the specified path
setwd("C:/Users/xpoumi/OneDrive/1. PhD GU/1. Research/Sina Copepodamide project/nMDS and Multivariate analysis/Sina_CA_analysis")

# Check if the 'pacman' package is installed, if not, install it
if (!require(pacman)) {
  install.packages("pacman")
}

# Load/install packages from GitHub
pacman::p_load_gh("Russel88/MicEco")

# Load/install packages from CRAN
pacman::p_load(
  devtools, vegan, readr, ggpubr, rstatix, ggh4x, knitr,
  gtsummary, webshot2, BiodiversityR, tidyverse
)

# Write the bibliography of loaded packages to a file named 'packages.bib'
write_bib(.packages(), "packages.bib")
```

### Targeted copepodamide analysis

#### Data import and preparation

```
# Read data from a CSV file into 'Sina_data' dataframe
Sina_data <- 
  read_delim("Arnoldt_targeted_analysis_data.csv",
             delim = ";", 
             escape_double = FALSE, 
             # Define column types for specific columns
             col_types = 
               cols(Dry_weight_µg = col_number(), 
                    CA_per_copepod_mol = col_number(), 
                    CA_per_copepod_pmol = col_number(), 
                    `448.3_CA` = col_number(), 
                    `686.5_CA` = col_number(),
                    `706.6_CA` = col_number(), 
                    `708.6_CA` = col_number(), 
                    `712.5_CA` = col_number(), 
                    `732.5_CA` = col_number(), 
                    `734.5_CA` = col_number(), 
                    `758.6_CA` = col_number(), 
                    `450.3_dhCA` = col_number(), 
                    `660.5_dhCA` = col_number(), 
                    `674.6_dhCA` = col_number(), 
                    `680.5_dhCA` = col_number(),
                    `686.5_dhCA` = col_number(), 
                    `688.6_dhCA` = col_number(), 
                    `706.5_dhCA` = col_number(),
                    `710.6_dhCA` = col_number(), 
                    `708.5_dhCA` = col_number(),
                    `712.6_dhCA` = col_number(), 
                    `714.6_dhCA` = col_number(),
                    `734.5_dhCA` = col_number(),
                    `736.6_dhCA` = col_number(),
                    `760.5_dhCA` = col_number()), 
             trim_ws = TRUE)


# Filter rows where the number of copepods is exactly 1
Sina_data <- Sina_data %>% filter(No_of_copepods == 1)

# Print the structure of the filtered Sina_data
str(Sina_data)
```

```
## spc_tbl_ [116 × 39] (S3: spec_tbl_df/tbl_df/tbl/data.frame)
##  $ Sample             : chr [1:116] "F1a" "F1b" "F1c" "F1d" ...
##  $ Type               : chr [1:116] "Freshwater" "Freshwater" "Freshwater" "Freshwater" ...
##  $ Site               : chr [1:116] "F1" "F1" "F1" "F1" ...
##  $ Time_diff_h        : num [1:116] 0.5 0.667 0.667 0.667 0.75 ...
##  $ Time_diff_min      : num [1:116] 30 40 40 40 45 55 55 65 65 65 ...
##  $ No_of_copepods     : num [1:116] 1 1 1 1 1 1 1 1 1 1 ...
##  $ Taxon              : chr [1:116] "Cyclops strenuus" "Cyclops strenuus" "Cyclops strenuus" "Cyclops strenuus" ...
##  $ Sex                : chr [1:116] "female" "female" "female" "female" ...
##  $ Stage              : chr [1:116] "adult" "adult" "adult" "adult" ...
##  $ Prosome_length_µm  : num [1:116] 920 1100 1000 820 1060 940 1040 1080 1120 1020 ...
##  $ Dry_weight_g       : num [1:116] 5.39e-06 8.37e-06 6.61e-06 4.06e-06 7.64e-06 5.68e-06 7.29e-06 8.00e-06 8.75e-06 6.95e-06 ...
##  $ Dry_weight_µg      : num [1:116] 5.39 8.37 6.61 4.06 7.64 5.68 7.29 8 8.75 6.94 ...
##  $ Dry_weight_ng      : num [1:116] 5385 8366 6614 4055 7636 ...
##  $ Data_file          : chr [1:116] "F1a.d" "F1b.d" "F1c.d" "F1d.d" ...
##  $ Acq_Date_Time      : POSIXct[1:116], format: "2022-02-03 18:28:00" "2022-02-03 18:54:00" ...
##  $ CA_per_copepod_mol : num [1:116] 4.00e-12 1.66e-11 1.84e-11 6.66e-12 1.87e-11 ...
##  $ CA_per_copepod_pmol: num [1:116] 4 16.6 18.36 6.66 18.72 ...
##  $ 448.3_CA           : num [1:116] 0 0 0 0 0 0 0 0 0 0 ...
##  $ 450.3_dhCA         : num [1:116] 0.00166 0.00459 0.00554 0.00515 0.00648 ...
##  $ 660.5_dhCA         : num [1:116] 0.00389 0.01523 0.00939 0.00636 0.02363 ...
##  $ 674.6_dhCA         : num [1:116] 0.00261 0.01855 0.0411 0.01095 0.03316 ...
##  $ 680.5_dhCA         : num [1:116] 0 0 0 0 0 0 0 0 0 0 ...
##  $ 686.5_dhCA         : num [1:116] 0.0365 0.0836 0.0904 0.0484 0.1182 ...
##  $ 686.5_CA           : num [1:116] 0 0 0 0 0 0 0 0 0 0 ...
##  $ 688.6_dhCA         : num [1:116] 0.00377 0.02141 0.02289 0.00662 0.02105 ...
##  $ 706.5_dhCA         : num [1:116] 0 0 0 0 0 0 0 0 0 0 ...
##  $ 706.6_CA           : num [1:116] 0 0 0 0 0 0 0 0 0 0 ...
##  $ 708.5_dhCA         : num [1:116] 0.0015 0.00711 0.00378 0.00204 0.00803 ...
##  $ 708.6_CA           : num [1:116] 0 0 0 0 0 0 0 0 0 0 ...
##  $ 710.6_dhCA         : num [1:116] 0.00835 0.07075 0.05861 0.01641 0.04762 ...
##  $ 712.5_CA           : num [1:116] 0 0 0 0 0 0 0 0 0 0 ...
##  $ 712.6_dhCA         : num [1:116] 0.00399 0.04105 0.05015 0.01111 0.03397 ...
##  $ 714.6_dhCA         : num [1:116] 0.011 0.0312 0.0455 0.0128 0.0437 ...
##  $ 732.5_CA           : num [1:116] 0 0 0 0 0 0 0 0 0 0 ...
##  $ 734.5_dhCA         : num [1:116] 0.0047 0.02474 0.0252 0.00899 0.02502 ...
##  $ 734.5_CA           : num [1:116] 0 0 0 0 0 0 0 0 0 0 ...
##  $ 736.6_dhCA         : num [1:116] 0.00103 0.00792 0.01146 0.00355 0.00659 ...
##  $ 758.6_CA           : num [1:116] 0 0 0 0 0 0 0 0 0 0 ...
##  $ 760.5_dhCA         : num [1:116] 0.001 0.00575 0.00312 0.00077 0.00701 ...
##  - attr(*, "spec")=
##   .. cols(
##   ..   Sample = col_character(),
##   ..   Type = col_character(),
##   ..   Site = col_character(),
##   ..   Time_diff_h = col_double(),
##   ..   Time_diff_min = col_double(),
##   ..   No_of_copepods = col_double(),
##   ..   Taxon = col_character(),
##   ..   Sex = col_character(),
##   ..   Stage = col_character(),
##   ..   Prosome_length_µm = col_double(),
##   ..   Dry_weight_g = col_double(),
##   ..   Dry_weight_µg = col_number(),
##   ..   Dry_weight_ng = col_double(),
##   ..   Data_file = col_character(),
##   ..   Acq_Date_Time = col_datetime(format = ""),
##   ..   CA_per_copepod_mol = col_number(),
##   ..   CA_per_copepod_pmol = col_number(),
##   ..   `448.3_CA` = col_number(),
##   ..   `450.3_dhCA` = col_number(),
##   ..   `660.5_dhCA` = col_number(),
##   ..   `674.6_dhCA` = col_number(),
##   ..   `680.5_dhCA` = col_number(),
##   ..   `686.5_dhCA` = col_number(),
##   ..   `686.5_CA` = col_number(),
##   ..   `688.6_dhCA` = col_number(),
##   ..   `706.5_dhCA` = col_number(),
##   ..   `706.6_CA` = col_number(),
##   ..   `708.5_dhCA` = col_number(),
##   ..   `708.6_CA` = col_number(),
##   ..   `710.6_dhCA` = col_number(),
##   ..   `712.5_CA` = col_number(),
##   ..   `712.6_dhCA` = col_number(),
##   ..   `714.6_dhCA` = col_number(),
##   ..   `732.5_CA` = col_number(),
##   ..   `734.5_dhCA` = col_number(),
##   ..   `734.5_CA` = col_number(),
##   ..   `736.6_dhCA` = col_number(),
##   ..   `758.6_CA` = col_number(),
##   ..   `760.5_dhCA` = col_number()
##   .. )
##  - attr(*, "problems")=<externalptr>
```

```
# Attach the filtered data, this makes the column names 
# directly accessible without specifying the dataframe
attach(Sina_data)

# Identify and select specific columns for descriptive/environmental data
data_id_target <- Sina_data %>%
  select(Sample, Type, Site, Time_diff_h, Time_diff_min, 
         Taxon, Dry_weight_ng, CA_per_copepod_pmol) %>% 
  as.data.frame()

# Subset the dataframe to create data for ordination
data_nMDS_target <- Sina_data %>% 
  select(!Sample & !Type & !Site & !Time_diff_h & 
           !Time_diff_min & !Taxon & !Sex & !Stage & 
         !Prosome_length_µm & !Dry_weight_g & !Dry_weight_µg & 
         !Dry_weight_ng & !Data_file & !Acq_Date_Time & 
         !CA_per_copepod_pmol & !No_of_copepods & !CA_per_copepod_mol) %>% 
  as.data.frame()

# Calculate proportions for data_nMDS_target
data_nMDS_target_props <- data_nMDS_target/rowSums(data_nMDS_target) # as proportions

# Display the first few rows of data_id_target
head(data_id_target)
```

```
## # A tibble: 6 × 8
##   Sample Type       Site  Time_diff_h Time_diff_min Taxon          Dry_weight_ng
##   <chr>  <chr>      <chr>       <dbl>         <dbl> <chr>                  <dbl>
## 1 F1a    Freshwater F1          0.5              30 Cyclops stren…         5385.
## 2 F1b    Freshwater F1          0.667            40 Cyclops stren…         8366.
## 3 F1c    Freshwater F1          0.667            40 Cyclops stren…         6614 
## 4 F1d    Freshwater F1          0.667            40 Cyclops stren…         4055.
## 5 F1e    Freshwater F1          0.75             45 Cyclops stren…         7636.
## 6 F1f    Freshwater F1          0.917            55 Cyclops stren…         5678.
## # ℹ 1 more variable: CA_per_copepod_pmol <dbl>
```

```
# Display the last few rows of data_id_target
tail(data_id_target)
```

```
## # A tibble: 6 × 8
##   Sample Type   Site  Time_diff_h Time_diff_min Taxon              Dry_weight_ng
##   <chr>  <chr>  <chr>       <dbl>         <dbl> <chr>                      <dbl>
## 1 M4h    Marine M4           2.08           125 Calanus sp               335575.
## 2 M4i    Marine M4           2.08           125 Unknown                  141286.
## 3 M4j    Marine M4           2.08           125 Temora longicornis         9617.
## 4 M4w    Marine M4           2.67           160 Centropages hamat…        16978.
## 5 M4y    Marine M4           2.25           135 Temora longicornis        25067.
## 6 M4z    Marine M4           2.67           160 Calanus sp               344887.
## # ℹ 1 more variable: CA_per_copepod_pmol <dbl>
```

```
# Display the first few rows of data_nMDS_target
head(data_nMDS_target)
```

```
## # A tibble: 6 × 22
##   `448.3_CA` `450.3_dhCA` `660.5_dhCA` `674.6_dhCA` `680.5_dhCA` `686.5_dhCA`
##        <dbl>        <dbl>        <dbl>        <dbl>        <dbl>        <dbl>
## 1          0      0.00166      0.00389      0.00261            0       0.0365
## 2          0      0.00459      0.0152       0.0185             0       0.0836
## 3          0      0.00554      0.00939      0.0411             0       0.0904
## 4          0      0.00515      0.00636      0.0110             0       0.0484
## 5          0      0.00648      0.0236       0.0332             0       0.118 
## 6          0      0.00739      0.0196       0.0317             0       0.112 
## # ℹ 16 more variables: `686.5_CA` <dbl>, `688.6_dhCA` <dbl>,
## #   `706.5_dhCA` <dbl>, `706.6_CA` <dbl>, `708.5_dhCA` <dbl>, `708.6_CA` <dbl>,
## #   `710.6_dhCA` <dbl>, `712.5_CA` <dbl>, `712.6_dhCA` <dbl>,
## #   `714.6_dhCA` <dbl>, `732.5_CA` <dbl>, `734.5_dhCA` <dbl>, `734.5_CA` <dbl>,
## #   `736.6_dhCA` <dbl>, `758.6_CA` <dbl>, `760.5_dhCA` <dbl>
```

```
# Display the first few rows of data_nMDS_target_props
head(data_nMDS_target_props)
```

```
## # A tibble: 6 × 22
##   `448.3_CA` `450.3_dhCA` `660.5_dhCA` `674.6_dhCA` `680.5_dhCA` `686.5_dhCA`
##        <dbl>        <dbl>        <dbl>        <dbl>        <dbl>        <dbl>
## 1          0       0.0207       0.0486       0.0326            0        0.456
## 2          0       0.0138       0.0459       0.0559            0        0.252
## 3          0       0.0151       0.0256       0.112             0        0.246
## 4          0       0.0387       0.0478       0.0823            0        0.363
## 5          0       0.0173       0.0631       0.0886            0        0.316
## 6          0       0.0217       0.0576       0.0931            0        0.328
## # ℹ 16 more variables: `686.5_CA` <dbl>, `688.6_dhCA` <dbl>,
## #   `706.5_dhCA` <dbl>, `706.6_CA` <dbl>, `708.5_dhCA` <dbl>, `708.6_CA` <dbl>,
## #   `710.6_dhCA` <dbl>, `712.5_CA` <dbl>, `712.6_dhCA` <dbl>,
## #   `714.6_dhCA` <dbl>, `732.5_CA` <dbl>, `734.5_dhCA` <dbl>, `734.5_CA` <dbl>,
## #   `736.6_dhCA` <dbl>, `758.6_CA` <dbl>, `760.5_dhCA` <dbl>
```

#### Ordination (nMDS)

##### Using concentration data

```
# Set a seed for reproducibility
set.seed(42)

# Perform Non-metric Multidimensional Scaling (nMDS) using the metaMDS function
nMDS_target <- metaMDS(data_nMDS_target,
                       distance = "bray",  # Bray-Curtis dissimilarity used as distance metric
                       k = 2,              # Number of dimensions to retain
                       try = 99,           # Number of random starts to attempt
                       trymax = 200,       # Maximum number of attempts
                       autotransform = FALSE) # Don't automatically transform the data
```

```
# Show the resulting ordination results
nMDS_target
```

```
## 
## Call:
## metaMDS(comm = data_nMDS_target, distance = "bray", k = 2, try = 99,      trymax = 200, autotransform = FALSE) 
## 
## global Multidimensional Scaling using monoMDS
## 
## Data:     data_nMDS_target 
## Distance: bray 
## 
## Dimensions: 2 
## Stress:     0.0756139 
## Stress type 1, weak ties
## Best solution was repeated 31 times in 99 tries
## The best solution was from try 46 (random start)
## Scaling: centring, PC rotation, halfchange scaling 
## Species: expanded scores based on 'data_nMDS_target'
```

##### Using proportions of total concentration data

```
# Set the seed for reproducibility in random processes
set.seed(42)

# Perform non-metric multidimensional scaling (nMDS) analysis on data_nMDS_target_props
nMDS_target_props <- metaMDS(
  data_nMDS_target_props,  # The data to be analyzed
  distance = "bray",       # The distance measure to be used
  k = 2,                   # The number of dimensions for the output
  try = 99,                # Number of iterations to attempt
  trymax = 200,            # Maximum number of iterations to attempt
  autotransform = FALSE   # Whether to automatically transform the data
)
```

```
# Show the resulting ordination results
nMDS_target_props
```

```
## 
## Call:
## metaMDS(comm = data_nMDS_target_props, distance = "bray", k = 2,      try = 99, trymax = 200, autotransform = FALSE) 
## 
## global Multidimensional Scaling using monoMDS
## 
## Data:     data_nMDS_target_props 
## Distance: bray 
## 
## Dimensions: 2 
## Stress:     0.1052285 
## Stress type 1, weak ties
## Best solution was repeated 2 times in 99 tries
## The best solution was from try 28 (random start)
## Scaling: centring, PC rotation, halfchange scaling 
## Species: expanded scores based on 'data_nMDS_target_props'
```

#### Plotting the nMDS results

##### Using nMDS on concentrations

```
# Extracting axis scores
datascores_target <- 
  as.data.frame(scores(nMDS_target, 
                       display = "sites")) # Extracts site scores and converts to a data frame

scores_target <- 
  cbind(data_id_target, 
        datascores_target)  # Combines data frames by columns

scores_target <-  
  scores_target %>%   # Selects specific columns for the final data frame
  select(Sample, Type, Site, NMDS1, NMDS2)


# Calculating centroids for Site
centroids_site_target <- # Computes mean values for each Site
  aggregate(cbind(NMDS1, NMDS2) ~ Site, 
            data = scores_target, 
            FUN = mean)  

centroids_type_target <- # Computes mean values for each Type
  aggregate(cbind(NMDS1, NMDS2) ~ Type, 
            data = scores_target, 
            FUN = mean)  


# Define colors for plotting
colours <- c("#005F07","#00B60D", "#00FD12",
             "#86AD6C", "#008080", "#99E56A", "#228B22",  # Greens
             "#0E1DFF", "#2472FA", "#5713FF", 
             "#657BC5", "#0000D1")  # Blues/Purples


# Create the nMDS plot
scores_target %>% 
  ggplot(aes(x = NMDS1, y = NMDS2)) +  # Define axes for the plot
  
  # Adding ellipses (95% confidence intervals)
  stat_ellipse(aes(fill = Type),  # Draws ellipses around points, colored by 'Type'
               geom = "polygon",  # Specifies the geometric object as a polygon
               alpha = 0.07,  # Sets the transparency of ellipses
               linewidth = 0.5,  # Sets the width of ellipse lines
               level = 0.95,  # Sets the confidence level for ellipses
               show.legend = FALSE) +  # Hides legend for this layer
               
  # Adding points with Site as fill color
  geom_point(aes(fill = Site),  # Adds points, colored by 'Site'
             colour = "black",  # Sets point border color
             size = 1.5,  # Sets point size
             shape = 21,  # Sets point shape (a circle with border)
             alpha = 0.3,  # Sets point transparency
             show.legend = FALSE) +  # Hides legend for this layer
  
  # Adding centroids with Site as fill color
  geom_point(aes(fill = Site),  # Adds centroids, colored by 'Site'
             data = centroids_site_target,  # Uses centroids data
             size = 2.5,  # Sets centroid size
             shape = 21,  # Sets centroid shape (a circle with border)
             alpha = 0.7,  # Sets centroid transparency
             colour = "black",  # Sets centroid border color
             show.legend = TRUE) +  # Shows legend for this layer

  # Applying classic theme
  theme_classic() +  # Applies a classic theme to the plot
  
  # Setting y-axis limits and expansion
  scale_y_continuous(limits = c(-2, 2.1),  # Sets y-axis limits
                     expand = c(0,0)) +  # Adjusts axis expansion
  
  # Setting x-axis limits, expansion, and breaks
  scale_x_continuous(limits = c(-4.5, 4),  # Sets x-axis limits
                     expand = c(0,0),  # Adjusts axis expansion
                     breaks = c(-4, -2, 0, 2, 4)) +  # Sets custom breaks on x-axis
  
  # Setting manual fill colors for Site
  scale_fill_manual(values = colours,   # Sets fill colors manually
                    name = "Site",  # Sets the legend title
                    labels = c("F1", "F2", "F3", 
                               "F4", "F5", "F6", 
                               "Fre", "M1", "M2", 
                               "M3", "M4", "Mar")) +  # Sets legend labels
  
  # Setting manual color values for Type
  scale_colour_manual(values = c("#228B22", "#0000D1")) +  # Sets color manually for Type

  # Setting theme parameters
  theme(
    plot.margin = unit(c(0.3, 0.2, 0.1, 0.1), "cm"),  # Sets plot margins
    
    axis.title.y = element_text(vjust = 0, 
                                size = 8),  # Sets y-axis title appearance
    
    axis.title.x = element_text(size = 8),  # Sets x-axis title appearance
    
    axis.text.y  = element_text(colour = "black", 
                                size = 8),  # Sets y-axis text appearance
    
    axis.text.x  = element_text(colour = "black", 
                                size = 8),  # Sets x-axis text appearance
    
    legend.text = element_text(colour = "black", 
                               size = 7),  # Sets legend text appearance
    
    legend.title = element_text(colour = "black", 
                                size = 8),  # Sets legend title appearance
    
    legend.position = c(0.92, 0.4),  # Sets legend position
    
    legend.margin = margin(unit(c(0, -22, 0, -20), "cm")),  # Sets legend margin
    
    legend.key.size = unit(0.3, 'cm')  # Sets legend key size
  )
```

```
# Saving the plot as a TIFF file
## ggsave("targeted_nMDS_new.tiff", width = 80, height = 65, units = "mm", dpi=700)
```

##### Using nMDS on proportions of total

```
# Extracting axis scores
datascores_target_props <- 
  as.data.frame(scores(nMDS_target_props, 
                       display = "sites")) # Extracts site scores and converts to a data frame

scores_target_props <- 
  cbind(data_id_target, 
        datascores_target_props)  # Combines data frames by columns

scores_target_props <-  
  scores_target_props %>%   # Selects specific columns for the final data frame
  select(Sample, Type, Site, NMDS1, NMDS2)


# Calculating centroids for Site
centroids_site_target_props <- # Computes mean values for each Site
  aggregate(cbind(NMDS1, NMDS2) ~ Site, 
            data = scores_target_props, 
            FUN = mean)  

centroids_type_target_props <- # Computes mean values for each Type
  aggregate(cbind(NMDS1, NMDS2) ~ Type, 
            data = scores_target_props, 
            FUN = mean)  


# Define colors for plotting
colours <- c("#005F07","#00B60D", "#00FD12",
             "#86AD6C", "#008080", "#99E56A", "#228B22",  # Greens
             "#0E1DFF", "#2472FA", "#5713FF", 
             "#657BC5", "#0000D1")  # Blues/Purples


# Create the nMDS plot
scores_target_props %>% 
  ggplot(aes(x = NMDS1, y = NMDS2)) +  # Define axes for the plot
  
  # Adding ellipses (95% confidence intervals)
  stat_ellipse(aes(fill = Type),  # Draws ellipses around points, colored by 'Type'
               geom = "polygon",  # Specifies the geometric object as a polygon
               alpha = 0.07,  # Sets the transparency of ellipses
               linewidth = 0.5,  # Sets the width of ellipse lines
               level = 0.95,  # Sets the confidence level for ellipses
               show.legend = FALSE) +  # Hides legend for this layer
               
  # Adding points with Site as fill color
  geom_point(aes(fill = Site),  # Adds points, colored by 'Site'
             colour = "black",  # Sets point border color
             size = 1.5,  # Sets point size
             shape = 21,  # Sets point shape (a circle with border)
             alpha = 0.3,  # Sets point transparency
             show.legend = FALSE) +  # Hides legend for this layer
  
  # Adding centroids with Site as fill color
  geom_point(aes(fill = Site),  # Adds centroids, colored by 'Site'
             data = centroids_site_target_props,  # Uses centroids data
             size = 2.5,  # Sets centroid size
             shape = 21,  # Sets centroid shape (a circle with border)
             alpha = 0.7,  # Sets centroid transparency
             colour = "black",  # Sets centroid border color
             show.legend = TRUE) +  # Shows legend for this layer

  # Applying classic theme
  theme_classic() +  # Applies a classic theme to the plot
  
  # Setting y-axis limits and expansion
  scale_y_continuous(limits = c(-1.5, 1.5),  # Sets y-axis limits
                     expand = c(0,0),       # Adjusts axis expansion
                     breaks = c(-1.5, 0, 1 )) +  # Sets custom breaks on y-axis
  
  # Setting x-axis limits, expansion, and breaks
  scale_x_continuous(limits = c(-2, 2),  # Sets x-axis limits
                     expand = c(0,0),  # Adjusts axis expansion
                     breaks = c(-4, -2, 0, 2, 4)) +  # Sets custom breaks on x-axis
  
  # Setting manual fill colors for Site
  scale_fill_manual(values = colours,   # Sets fill colors manually
                    name = "Site",  # Sets the legend title
                    labels = c("F1", "F2", "F3", 
                               "F4", "F5", "F6", 
                               "Fre", "M1", "M2", 
                               "M3", "M4", "Mar")) +  # Sets legend labels
  
  # Setting manual color values for Type
  scale_colour_manual(values = c("#228B22", "#0000D1")) +  # Sets color manually for Type

  # Setting theme parameters
  theme(
    plot.margin = unit(c(0.3, 0.2, 0.1, 0.1), "cm"),  # Sets plot margins
    
    axis.title.y = element_text(vjust = 0, 
                                size = 8),  # Sets y-axis title appearance
    
    axis.title.x = element_text(size = 8),  # Sets x-axis title appearance
    
    axis.text.y  = element_text(colour = "black", 
                                size = 8),  # Sets y-axis text appearance
    
    axis.text.x  = element_text(colour = "black", 
                                size = 8),  # Sets x-axis text appearance
    
    legend.text = element_text(colour = "black", 
                               size = 7),  # Sets legend text appearance
    
    legend.title = element_text(colour = "black", 
                                size = 8),  # Sets legend title appearance
    
    legend.position = c(0.1, 0.4),  # Sets legend position
    
    legend.margin = margin(unit(c(0, -22, 0, -20), "cm")),  # Sets legend margin
    
    legend.key.size = unit(0.3, 'cm')  # Sets legend key size
  )
```

```
# Saving the plot as a TIFF file
## ggsave("targeted_nMDS_props.tiff", width = 80, height = 65, units = "mm", dpi=700)
```

#### Multivariate analysis - PERMANOVA

##### On concentration data

```
# Testing for homogeneity of dispersion
set.seed(42)  # Set the random seed for reproducibility

targeted_dist <- 
  vegdist(data_nMDS_target,
          method = "bray")  # Calculate Bray-Curtis dissimilarity

dispersion_targeted <- # Compute multivariate homogeneity of group dispersions
  betadisper(targeted_dist, 
             group = data_id_target$Type)  

dispersion_targeted  # Display the results
```

```
## 
##  Homogeneity of multivariate dispersions
## 
## Call: betadisper(d = targeted_dist, group = data_id_target$Type)
## 
## No. of Positive Eigenvalues: 66
## No. of Negative Eigenvalues: 49
## 
## Average distance to median:
## Freshwater     Marine 
##     0.4802     0.5076 
## 
## Eigenvalues for PCoA axes:
## (Showing 8 of 115 eigenvalues)
##  PCoA1  PCoA2  PCoA3  PCoA4  PCoA5  PCoA6  PCoA7  PCoA8 
## 10.009  6.834  4.051  3.546  2.140  1.370  1.284  1.095
```

```
# Plotting the homogeneity of dispersion
plot(dispersion_targeted)  # Create a plot to visualize the dispersion
```

```
# Perform an ANOVA on the dispersion
anova(dispersion_targeted)  # Tests for significant differences in group dispersions
```

```
## # A tibble: 2 × 5
##      Df `Sum Sq` `Mean Sq` `F value` `Pr(>F)`
##   <int>    <dbl>     <dbl>     <dbl>    <dbl>
## 1     1   0.0219    0.0219      1.54    0.217
## 2   114   1.62      0.0142     NA      NA
```

```
# Permutation test for homogeneity of dispersion
permutest(dispersion_targeted, permutations = 9999)
```

```
## 
## Permutation test for homogeneity of multivariate dispersions
## Permutation: free
## Number of permutations: 9999
## 
## Response: Distances
##            Df  Sum Sq  Mean Sq      F N.Perm Pr(>F)
## Groups      1 0.02189 0.021886 1.5404   9999 0.2136
## Residuals 114 1.61966 0.014208
```

```
# Performs a permutation test to assess if observed dispersion differences are significant

# Post-hoc test for differences in group means (Tukey's HSD)
# If ANOVA indicates significant differences in dispersions, this test identifies specific pairs of groups that differ
TukeyHSD(dispersion_targeted)
```

```
##   Tukey multiple comparisons of means
##     95% family-wise confidence level
## 
## Fit: aov(formula = distances ~ group, data = df)
## 
## $group
##                         diff         lwr        upr     p adj
## Marine-Freshwater 0.02748784 -0.01638566 0.07136133 0.2171015
```

```
# Testing for differences in centroid position between the habitats (type)
fit_targeted <- adonis2(data_nMDS_target ~ Type, 
                        data=data_id_target, 
                        permutations=9999,
                        method="bray", 
                        by = "margin")

fit_targeted  # Display the results of the adonis test
```

```
## # A tibble: 3 × 5
##      Df SumOfSqs    R2     F `Pr(>F)`
##   <dbl>    <dbl> <dbl> <dbl>    <dbl>
## 1     1     6.28 0.175  24.1   0.0001
## 2   114    29.7  0.825  NA    NA     
## 3   115    35.9  1      NA    NA
```

```
 # Compute partial Omega-squared for the adonis model predictors
adonis_OmegaSq(fit_targeted, partial = TRUE)
```

```
## # A tibble: 3 × 5
##      Df SumOfSqs     F parOmegaSq `Pr(>F)`
##   <dbl>    <dbl> <dbl>      <dbl>    <dbl>
## 1     1     6.28  24.1      0.166   0.0001
## 2   114    29.7   NA       NA      NA     
## 3   115    35.9   NA       NA      NA
```

```
# Similar adonis tests with taxon added as predictor
fit2 <- adonis2(data_nMDS_target ~ Type + Taxon, 
                data=data_id_target, 
                permutations=9999, 
                method="bray", 
                by = "margin")

fit2  # Display the results
```

```
## # A tibble: 4 × 5
##      Df SumOfSqs     R2     F `Pr(>F)`
##   <dbl>    <dbl>  <dbl> <dbl>    <dbl>
## 1     1     1.39 0.0388  7.75   0.0001
## 2    12    11.3  0.315   5.26   0.0001
## 3   102    18.3  0.510  NA     NA     
## 4   115    35.9  1      NA     NA
```

```
adonis_OmegaSq(fit2, partial = TRUE)  # Compute partial Omega-squared
```

```
## # A tibble: 4 × 5
##      Df SumOfSqs     F parOmegaSq `Pr(>F)`
##   <dbl>    <dbl> <dbl>      <dbl>    <dbl>
## 1     1     1.39  7.75     0.0550   0.0001
## 2    12    11.3   5.26     0.306    0.0001
## 3   102    18.3  NA       NA       NA     
## 4   115    35.9  NA       NA       NA
```

```
# SIMPER analysis to identify important variables driving dissimilarity
# SIMPER helps identify which variables contribute the most to dissimilarity between groups
simper_targeted <- simper(data_nMDS_target, data_id_target$Type, 
                          permutations = 9999)

summary(simper_targeted, ordered = TRUE)  # Summarize the SIMPER results
```

```
## 
## Contrast: Freshwater_Marine 
## 
##             average       sd    ratio      ava      avb cumsum      p    
## 708.5_dhCA 0.137450 0.123420 1.113700 0.007997 0.036190  0.163 0.0001 ***
## 732.5_CA   0.068650 0.103170 0.665400 0.000008 0.022290  0.244 0.0001 ***
## 686.5_dhCA 0.065140 0.076260 0.854100 0.017859 0.001090  0.321 0.5460    
## 712.6_dhCA 0.058030 0.058660 0.989400 0.013336 0.000000  0.390 0.0008 ***
## 734.5_dhCA 0.057010 0.039620 1.439200 0.007936 0.014540  0.458 0.0004 ***
## 710.6_dhCA 0.055450 0.046370 1.195900 0.014159 0.005380  0.523 1.0000    
## 706.5_dhCA 0.046810 0.068560 0.682700 0.000002 0.013770  0.579 0.0001 ***
## 448.3_CA   0.046640 0.085660 0.544400 0.000000 0.016370  0.634 0.0001 ***
## 706.6_CA   0.045970 0.076270 0.602700 0.000000 0.009600  0.689 0.0001 ***
## 688.6_dhCA 0.042940 0.047260 0.908700 0.005294 0.008990  0.739 0.0032 ** 
## 714.6_dhCA 0.038410 0.037290 1.030000 0.009578 0.001100  0.785 0.0037 ** 
## 660.5_dhCA 0.035430 0.031510 1.124400 0.007041 0.004610  0.827 1.0000    
## 760.5_dhCA 0.029980 0.030970 0.968100 0.001944 0.008310  0.862 0.0001 ***
## 758.6_CA   0.028650 0.047720 0.600500 0.000010 0.012210  0.896 0.0001 ***
## 450.3_dhCA 0.026440 0.041430 0.638100 0.001556 0.004490  0.928 0.0001 ***
## 674.6_dhCA 0.016370 0.020870 0.784400 0.004911 0.000000  0.947 0.6728    
## 736.6_dhCA 0.011500 0.012110 0.949400 0.002755 0.000000  0.961 0.0021 ** 
## 680.5_dhCA 0.011180 0.024540 0.455500 0.000000 0.003510  0.974 0.0001 ***
## 686.5_CA   0.010480 0.022130 0.473500 0.000000 0.004690  0.986 0.0001 ***
## 708.6_CA   0.009530 0.014930 0.638600 0.000000 0.002150  0.998 0.0001 ***
## 734.5_CA   0.001910 0.004120 0.462700 0.000267 0.000000  1.000 1.0000    
## 712.5_CA   0.000080 0.000250 0.337100 0.000010 0.000000  1.000 0.9992    
## ---
## Signif. codes:  0 '***' 0.001 '**' 0.01 '*' 0.05 '.' 0.1 ' ' 1
## Permutation: free
## Number of permutations: 9999
```

##### On proportions of total concentration

```
# Testing for homogeneity of dispersion
set.seed(42)  # Set the random seed for reproducibility

targeted_dist_props <- 
  vegdist(data_nMDS_target_props,
          method = "bray")  # Calculate Bray-Curtis dissimilarity

dispersion_targeted_props <- # Compute multivariate homogeneity of group dispersions
  betadisper(targeted_dist_props, 
             group = data_id_target$Type)  

dispersion_targeted_props  # Display the results
```

```
## 
##  Homogeneity of multivariate dispersions
## 
## Call: betadisper(d = targeted_dist_props, group = data_id_target$Type)
## 
## No. of Positive Eigenvalues: 52
## No. of Negative Eigenvalues: 63
## 
## Average distance to median:
## Freshwater     Marine 
##     0.2311     0.3947 
## 
## Eigenvalues for PCoA axes:
## (Showing 8 of 115 eigenvalues)
##   PCoA1   PCoA2   PCoA3   PCoA4   PCoA5   PCoA6   PCoA7   PCoA8 
## 10.6314  5.7014  1.8339  0.8523  0.7614  0.6976  0.6194  0.4865
```

```
# Plotting the homogeneity of dispersion
plot(dispersion_targeted_props)  # Create a plot to visualize the dispersion
```

```
# Perform an ANOVA on the dispersion
anova(dispersion_targeted_props)  # Tests for significant differences in group dispersions
```

```
## # A tibble: 2 × 5
##      Df `Sum Sq` `Mean Sq` `F value`  `Pr(>F)`
##   <int>    <dbl>     <dbl>     <dbl>     <dbl>
## 1     1    0.775   0.775        81.6  4.95e-15
## 2   114    1.08    0.00949      NA   NA
```

```
# Permutation test for homogeneity of dispersion
permutest(dispersion_targeted_props, permutations = 9999)
```

```
## 
## Permutation test for homogeneity of multivariate dispersions
## Permutation: free
## Number of permutations: 9999
## 
## Response: Distances
##            Df  Sum Sq Mean Sq      F N.Perm Pr(>F)    
## Groups      1 0.77457 0.77457 81.587   9999  1e-04 ***
## Residuals 114 1.08229 0.00949                         
## ---
## Signif. codes:  0 '***' 0.001 '**' 0.01 '*' 0.05 '.' 0.1 ' ' 1
```

```
# Post-hoc test for differences in group means (Tukey's HSD)
# If ANOVA indicates significant differences in dispersions, this test identifies specific pairs of groups that differ
TukeyHSD(dispersion_targeted_props)
```

```
##   Tukey multiple comparisons of means
##     95% family-wise confidence level
## 
## Fit: aov(formula = distances ~ group, data = df)
## 
## $group
##                        diff       lwr      upr p adj
## Marine-Freshwater 0.1635267 0.1276624 0.199391     0
```

```
# Testing for differences in centroid position between the habitats (type)
fit_targeted_props <- adonis2(data_nMDS_target_props ~ Type, 
                        data=data_id_target, 
                        permutations=9999,
                        method="bray", 
                        by = "margin")

fit_targeted_props  # Display the results of the adonis test
```

```
## # A tibble: 3 × 5
##      Df SumOfSqs    R2     F `Pr(>F)`
##   <dbl>    <dbl> <dbl> <dbl>    <dbl>
## 1     1     9.39 0.424  84.0   0.0001
## 2   114    12.7  0.576  NA    NA     
## 3   115    22.1  1      NA    NA
```

```
adonis_OmegaSq(fit_targeted_props, partial = TRUE)  # Compute partial R-squared for the adonis test
```

```
## # A tibble: 3 × 5
##      Df SumOfSqs     F parOmegaSq `Pr(>F)`
##   <dbl>    <dbl> <dbl>      <dbl>    <dbl>
## 1     1     9.39  84.0      0.417   0.0001
## 2   114    12.7   NA       NA      NA     
## 3   115    22.1   NA       NA      NA
```

```
# Similar adonis tests with taxon added as predictor
fit2_props <- adonis2(data_nMDS_target_props ~ Type + Taxon, 
                data=data_id_target, 
                permutations=9999, 
                method="bray", 
                by = "margin")

fit2_props  # Display the results
```

```
## # A tibble: 4 × 5
##      Df SumOfSqs     R2     F `Pr(>F)`
##   <dbl>    <dbl>  <dbl> <dbl>    <dbl>
## 1     1     1.53 0.0693 26.7    0.0001
## 2    12     6.87 0.310   9.95   0.0001
## 3   102     5.87 0.265  NA     NA     
## 4   115    22.1  1      NA     NA
```

```
adonis_OmegaSq(fit2_props, partial = TRUE)  # Compute partial R-squared
```

```
## # A tibble: 4 × 5
##      Df SumOfSqs     F parOmegaSq `Pr(>F)`
##   <dbl>    <dbl> <dbl>      <dbl>    <dbl>
## 1     1     1.53 26.7       0.181   0.0001
## 2    12     6.87  9.95      0.481   0.0001
## 3   102     5.87 NA        NA      NA     
## 4   115    22.1  NA        NA      NA
```

```
# SIMPER analysis to identify important variables driving dissimilarity
simper_targeted_props <- simper(data_nMDS_target_props, data_id_target$Type, 
                          permutations = 9999)
summary(simper_targeted_props, ordered = TRUE)  # Summarize the SIMPER results
```

```
## 
## Contrast: Freshwater_Marine 
## 
##            average      sd   ratio     ava     avb cumsum      p    
## 708.5_dhCA 0.09211 0.06725 1.36960 0.06743 0.22555  0.126 0.0001 ***
## 712.6_dhCA 0.07967 0.04189 1.90190 0.15934 0.00000  0.236 0.0001 ***
## 686.5_dhCA 0.06868 0.04697 1.46220 0.14346 0.00776  0.330 0.0001 ***
## 732.5_CA   0.05181 0.06624 0.78220 0.00048 0.10404  0.402 0.0001 ***
## 714.6_dhCA 0.05101 0.02445 2.08650 0.10581 0.00379  0.472 0.0001 ***
## 710.6_dhCA 0.05089 0.02614 1.94680 0.13805 0.03910  0.542 0.0001 ***
## 688.6_dhCA 0.04158 0.03490 1.19140 0.09049 0.07547  0.599 0.0119 *  
## 706.6_CA   0.03785 0.05409 0.69980 0.00000 0.07570  0.651 0.0001 ***
## 448.3_CA   0.03676 0.05343 0.68810 0.00000 0.07353  0.701 0.0001 ***
## 706.5_dhCA 0.03592 0.04284 0.83840 0.00028 0.07209  0.750 0.0001 ***
## 660.5_dhCA 0.03099 0.02342 1.32340 0.07905 0.02837  0.793 0.0001 ***
## 760.5_dhCA 0.02427 0.02401 1.01100 0.03621 0.05532  0.827 0.0002 ***
## 758.6_CA   0.02208 0.02833 0.77950 0.00152 0.04457  0.857 0.0001 ***
## 450.3_dhCA 0.02203 0.02884 0.76390 0.02284 0.04805  0.887 0.0034 ** 
## 734.5_dhCA 0.02194 0.01498 1.46450 0.08339 0.09782  0.917 0.0013 ** 
## 674.6_dhCA 0.01752 0.01348 1.29950 0.03503 0.00000  0.941 0.0001 ***
## 736.6_dhCA 0.01469 0.00821 1.78860 0.02938 0.00000  0.962 0.0001 ***
## 686.5_CA   0.00844 0.01400 0.60310 0.00000 0.01689  0.973 0.0001 ***
## 708.6_CA   0.00806 0.01047 0.76980 0.00000 0.01612  0.984 0.0001 ***
## 680.5_dhCA 0.00792 0.01592 0.49720 0.00000 0.01583  0.995 0.0001 ***
## 734.5_CA   0.00311 0.00485 0.64010 0.00621 0.00000  0.999 0.0001 ***
## 712.5_CA   0.00051 0.00279 0.18200 0.00102 0.00000  1.000 0.4800    
## ---
## Signif. codes:  0 '***' 0.001 '**' 0.01 '*' 0.05 '.' 0.1 ' ' 1
## Permutation: free
## Number of permutations: 9999
```

```
# SIMPER helps identify which variables contribute the most to dissimilarity between groups
```

There are no significant differences in the results due to using
concentrations vs proportions of concentrations, we will therefore use
the proportions to homogenise with the precursor analysis further down
the pipeline (which only has proportional data).

#### Testing for the effects of copepod size on copepodamide content

##### ANCOVA

After some exploratory analysis, we found that log-transformation of
both continuous variables yielded the most normally distributed
residuals. We will therefore apply the natural log-transformation for
the final analysis.

###### Fitting the model

```
# Convert 'Type' and 'Site' columns to factor variables
Sina_data$Type <- as_factor(Sina_data$Type)  
Sina_data$Site <- as_factor(Sina_data$Site)  

# Perform an ANCOVA (Analysis of Covariance) model
targeted_ancova <- lm(log(CA_per_copepod_pmol) ~ log(Dry_weight_µg) + Type, 
                data =  filter(Sina_data, Taxon != "Harpacticoid cf Microsetella"))
```

###### Checking assumptions for ANCOVA

Linearity: Here we examine if there is a linear relationship between
the covariate and the response variable.

```
# Define a custom color palette
colours <- c( "#002F07", "#0001F5")

# Filter out rows where 'Taxon' is not equal to "Harpacticoid cf Microsetella"
# and create a scatter plot with a linear regression line
Sina_data %>%  
  filter(Taxon != "Harpacticoid cf Microsetella") %>%  # Filter the data
  ggplot(aes(x = log(Dry_weight_µg),   # X-axis: log-transformed 'Dry_weight_µg'
             y = log(CA_per_copepod_pmol))) +  # Y-axis: log-transformed 'CA_per_copepod_pmol'

  # Add a linear regression line, colored by 'Type'
  geom_smooth(aes(colour = Type),   # Color the line by the 'Type' variable
              method = "lm",       # Fit a linear model for the line
              se = FALSE,          # Do not display standard error ribbon
              alpha = 0.2) +       # Set transparency of the line

  # Set manual color scale using predefined colors
  scale_color_manual(values =  colours) + 

  theme_classic()  # Use the classic ggplot2 theme
```

```
# Define a custom color palette with various shades of greens and blues
colours <- c("#005F07","#00B60D", "#00FD12","#86AD6C","#008080", "#99E56A", 
             "#6D26AB", "#B672F1", "#0056F5", "#0001F5") 

# Filter out rows where 'Taxon' is not equal to "Harpacticoid cf Microsetella"
# and create a scatter plot with a linear regression line
Sina_data %>%  
  filter(Taxon != "Harpacticoid cf Microsetella") %>%  # Filter the data
  ggplot(aes(x = log(Dry_weight_µg),   # X-axis: log-transformed 'Dry_weight_µg'
             y = log(CA_per_copepod_pmol))) +  # Y-axis: log-transformed 'CA_per_copepod_pmol'

  # Add a linear regression line, colored by 'Site'
  geom_smooth(aes(colour = Site),   # Color the line by the 'Site' variable
              method = "lm",       # Fit a linear model for the line
              se = FALSE,          # Do not display standard error ribbon
              alpha = 0.2) +       # Set transparency of the line

  # Set manual color scale using predefined colors
  scale_color_manual(values =  colours) + 

  theme_classic()  # Use the classic ggplot2 theme
```

Homogeneity of regression slopes: Here we test to see that there is
no interaction between the covariate and the grouping factors

```
# Filter out rows where 'Taxon' is not equal to "Harpacticoid cf Microsetella"
# and perform an ANOVA test for the specified model formula
Sina_data %>% 
  filter(Taxon != "Harpacticoid cf Microsetella") %>%  # Filter the data
  
  anova_test(
    log(CA_per_copepod_pmol) ~ log(Dry_weight_µg) + Type + log(Dry_weight_µg):Type)
```

```
## # A tibble: 3 × 7
##   Effect                    DFn   DFd     F     p `p<.05`   ges
##   <chr>                   <dbl> <dbl> <dbl> <dbl> <chr>   <dbl>
## 1 log(Dry_weight_µg)          1   111 2.76  0.1   ""      0.024
## 2 Type                        1   111 2.22  0.139 ""      0.02 
## 3 log(Dry_weight_µg):Type     1   111 0.929 0.337 ""      0.008
```

Result: no interaction.

Normality of residuals:

```
# Perform the Shapiro-Wilk test for normality on the residuals of the ANCOVA model
shapiro.test(targeted_ancova$residuals)
```

```
## 
##  Shapiro-Wilk normality test
## 
## data:  targeted_ancova$residuals
## W = 0.96466, p-value = 0.003961
```

```
# Create a density plot of the residuals of the ANCOVA model
plot(density(targeted_ancova$residuals))
```

Result: Residuals are NOT normally distributed

###### ANCOVA test

```
# Obtain a summary of the ANCOVA model
summary.lm(targeted_ancova)
```

```
## 
## Call:
## lm(formula = log(CA_per_copepod_pmol) ~ log(Dry_weight_µg) + 
##     Type, data = filter(Sina_data, Taxon != "Harpacticoid cf Microsetella"))
## 
## Residuals:
##     Min      1Q  Median      3Q     Max 
## -4.9445 -0.8417  0.1740  1.0214  2.5633 
## 
## Coefficients:
##                    Estimate Std. Error t value Pr(>|t|)  
## (Intercept)          0.2780     0.2751   1.011   0.3144  
## log(Dry_weight_µg)   0.1904     0.1146   1.661   0.0994 .
## TypeMarine           0.5241     0.3520   1.489   0.1392  
## ---
## Signif. codes:  0 '***' 0.001 '**' 0.01 '*' 0.05 '.' 0.1 ' ' 1
## 
## Residual standard error: 1.481 on 112 degrees of freedom
## Multiple R-squared:  0.1042, Adjusted R-squared:  0.08824 
## F-statistic: 6.516 on 2 and 112 DF,  p-value: 0.002103
```

```
# Detailed  output with effect sizes (partial eta-squared)
anova_test(targeted_ancova, detailed = TRUE, effect.size = "pes")
```

```
## # A tibble: 2 × 9
##   Effect               SSn   SSd   DFn   DFd     F     p `p<.05`   pes
##   <chr>              <dbl> <dbl> <dbl> <dbl> <dbl> <dbl> <chr>   <dbl>
## 1 log(Dry_weight_µg)  6.05  246.     1   112  2.76 0.099 ""      0.024
## 2 Type                4.86  246.     1   112  2.22 0.139 ""      0.019
```

##### The plot (copepodamide content as a function of dry mass)

```
# Define colours to use in the plot
colours <- c("#005F07","#00B60D", "#00FD12","#86AD6C",
             "#008080", "#99E56A", "#228B22", #Greens
              "#3A0CA3", "#3F37C9", "#0000D1" , "#4361EE", "#0218F5")## Blues

# Filter out specific Taxon and create a scatter plot
Sina_data %>% 
  filter(Taxon != "Harpacticoid cf Microsetella") %>% 
  ggplot(aes(x = log(Dry_weight_µg), 
             y = log(CA_per_copepod_pmol),
             colour = Site)) +
  
  # Add smoothed lines by Type with transparency and legend turned off
  geom_line(
    aes(colour = Type),  # Set line color by Type
    stat = "smooth",     # Add a smoothing line
    method = "lm",       # Use linear regression for smoothing
    linewidth = 1.5,     # Set the width of the line
    alpha = 0.5,         # Set transparency of the line
    show.legend = FALSE  # Don't include this in the legend
  )+
  
  # Add a shaded area around the smoothed lines with legend turned off
  geom_ribbon(
    stat='smooth',       # Add a smoothing line
    method = "lm",      # Use linear regression for smoothing
    se=TRUE,             # Include standard error
    alpha=0.1,           # Set transparency of the ribbon
    show.legend = FALSE, # Don't include this in the legend
    aes(color = NULL, group = Type) # Color and group aesthetics
  ) +
  
  # Add data points with some transparency and legend turned on
  geom_point(
    alpha = 0.1,         # Set transparency of the points
    show.legend = TRUE   # Include this in the legend
  )+

  # Add smoothed lines with lower transparency and no legend
  geom_line(
    stat = "smooth",     # Add a smoothing line
    method = "lm",       # Use linear regression for smoothing
    linewidth = 0.5,     # Set the width of the line
    alpha = 0.2,         # Set transparency of the line
    show.legend = FALSE  # Don't include this in the legend
  )+
  
  # Add a separate smoothed line with custom appearance and no legend
  geom_smooth(
    colour = "black",    # Set line color
    method = "lm",       # Use linear regression for smoothing
    se = FALSE,          # Don't include standard error
    linewidth = 1,       # Set the width of the line
    linetype = "dotted", # Set line type
    show.legend = FALSE  # Don't include this in the legend
  )+
  
  # Customize the x-axis label
  xlab(bquote(Ln~Dry~mass~(µg)))+
  
  # Customize the y-axis label
  ylab(bquote(Ln~Copepodamide~amount~(pmol~ind^-1)))+

  # Set custom colors and labels for Site
  scale_colour_manual(
    values = colours,   # Set custom colors
    labels = c("F1", "F2", "F3", "F4", "F5", "F6", "Fre", "M1", "M2", "M3", "M4", "Mar") # Set legend labels
  )+
  
  # Apply a classic theme with specified margins and text sizes
  theme_classic()+
  
  # Further customize plot appearance
  theme(
    plot.margin = unit(c(0.3, 0.5, 0.1, 0), "cm"), # Set plot margins
    axis.title.y = element_text(vjust = 0, size = 8), # Customize y-axis title
    axis.title.x = element_text(size = 8), # Customize x-axis title
    axis.text.y  = element_text(colour = "black", size = 8), # Customize y-axis text
    axis.text.x  = element_text(colour = "black", size = 8), # Customize x-axis text
    legend.text = element_text(colour = "black", size = 7), # Customize legend text
    legend.title = element_text(colour = "black", size = 8), # Customize legend title
    legend.position = c(0.95, 0.35), # Adjust legend position
    legend.background = element_rect(fill='transparent', colour = 'transparent'), # Transparent legend bg
    legend.box.background = element_rect(fill='transparent', colour = 'transparent'), # Transparent legend panel
    legend.key.size = unit(0.2, 'cm') # Control the size of the legend key
  ) +
  
  # Customize the appearance of color and size legends
  guides(
    colour = guide_legend(ncol = 1, override.aes = list(alpha = 0.1)), # Set color legend appearance
    size = guide_legend(override.aes = list(size = FALSE)) # Set size legend appearance
  )
```

```
# Save the plot as a PNG image with specified dimensions and resolution
## ggsave("CA_over_mass.png", width = 80, height = 70, units = "mm", dpi=700)
```

###### Same plot, without logarithmic scale

```
# Define colours to use in the plot
colours <- c("#005F07","#00B60D", "#00FD12",
             "#86AD6C", "#008080", "#99E56A", "#228B22", #Greens
             "#480CA8", "#3A0CA3", "#3F37C9", "#0218F5", "#4361EE", "#0000D1")## Blues

# Filter out specific Taxon and create a scatter plot
Sina_data %>%
  filter(Taxon != "Harpacticoid cf Microsetella") %>% 
  
  ggplot(aes(x = (Dry_weight_µg), 
             y = (CA_per_copepod_pmol),
             colour = Site)) +
  
  # Add data points with transparency and size
  geom_point(
    alpha = 0.2,   # Set transparency of the points
    size = 1,      # Set point size
    show.legend = FALSE  # Don't include this in the legend
  )+
  
  # Add a smoothed line with specified appearance and legend
  geom_line(
    stat = "smooth",   # Add a smoothing line
    method = "lm",     # Use linear regression for smoothing
    linewidth = 0.7,   # Set the width of the line
    alpha = 0.5,       # Set transparency of the line
    show.legend = TRUE # Include this in the legend
  )+
  
  # Add a smoothed line with specified appearance and no legend
  geom_smooth(
    aes(colour = Type),  # Set line color by Type
    method = "lm",       # Use linear regression for smoothing
    se = TRUE,           # Include standard error
    alpha = 0.2,         # Set transparency of the line
    linewidth = 1,       # Set the width of the line
    linetype = "solid",  # Set line type
    show.legend = FALSE  # Don't include this in the legend
  )+
  
  # Customize the x-axis label
  xlab(bquote(Dry~mass~(µg)))+
  
  # Customize the y-axis label
  ylab(bquote(Copepodamide~amount~(pmol~ind^-1)))+
  
  # Set custom colors for the color scale
  scale_colour_manual(values = colours) +
  
  # Apply a classic theme with specified margins and text sizes
  theme_classic()+
  
  # Further customize plot appearance
  theme(
    plot.margin = unit(c(0.3, 0.5, 0.1, 0), "cm"), # Set plot margins
    axis.title.y = element_text(vjust = 0, size = 8), # Customize y-axis title
    axis.title.x = element_text(size = 8), # Customize x-axis title
    axis.text.y  = element_text(colour = "black", size = 8), # Customize y-axis text
    axis.text.x  = element_text(colour = "black", size = 8), # Customize x-axis text
    legend.text = element_text(colour = "black", size = 7), # Customize legend text
    legend.title = element_text(colour = "black", size = 8), # Customize legend title
    legend.position = c(0.85, 0.21), # Adjust legend position
    legend.background = element_rect(fill='transparent', colour = 'transparent'), # Transparent legend bg
    legend.box.background = element_rect(fill='transparent', colour = 'transparent'), # Transparent legend panel
    legend.key.size = unit(0.35, 'cm') # Control the size of the legend key
  ) +
  
  # Customize the appearance of the legend
  guides(
    colour = guide_legend(
      ncol = 2, # Set number of columns in the legend
      override.aes = list(alpha = 1, linewidth =1.25) # Override legend aesthetics
    )
  )
```

```
# Save the plot as a TIFF image with specified dimensions and resolution
## ggsave("Copepodamide_over_mass_noLN.tiff", width = 75, height = 65, units = "mm", dpi=700)
```

Because there is no effect of type, and no interaction effect, we
will remove type as a predictor and obtain an overall linear
relationship between dry mass and copepodamide content.

##### Overall linear relationship between copepod dry mass and copepodamide content

###### Model

```
# Fitting a linear regression model
cop_reg_no_microsetella <- lm(log(CA_per_copepod_pmol) ~ log(Dry_weight_µg), 
                              data = filter(Sina_data, Dry_weight_ng > 1))

# Summary of the linear regression model
summary.lm(cop_reg_no_microsetella)
```

```
## 
## Call:
## lm(formula = log(CA_per_copepod_pmol) ~ log(Dry_weight_µg), 
##     data = filter(Sina_data, Dry_weight_ng > 1))
## 
## Residuals:
##     Min      1Q  Median      3Q     Max 
## -4.9509 -0.8940  0.1439  1.0620  2.8038 
## 
## Coefficients:
##                    Estimate Std. Error t value Pr(>|t|)   
## (Intercept)         0.25033    0.27595   0.907  0.36625   
## log(Dry_weight_µg)  0.29604    0.09051   3.271  0.00142 **
## ---
## Signif. codes:  0 '***' 0.001 '**' 0.01 '*' 0.05 '.' 0.1 ' ' 1
## 
## Residual standard error: 1.489 on 113 degrees of freedom
## Multiple R-squared:  0.0865, Adjusted R-squared:  0.07841 
## F-statistic:  10.7 on 1 and 113 DF,  p-value: 0.001421
```

Note that there is one outlier, the single Harpacticoid c.f
*Microsetella* with very small mass (0.12 ng, more than three
order of magnitude smaller than the next smallest, which must be an
error in the length-weight conversion), the exclusion of which does not
change either the slope or intercept, only the significance values for
the intercept and the slope. We will use the model that results in more
conservative values, which is the one excluding the harpacticoid
sample.

###### Plot of overall linear relationship

```
# Define a vector of custom colors
colours <- c("#005F07","#00B60D", "#00FD12",
             "#86AD6C", "#008080", "#99E56A", 
             "#480CA8", "#3A0CA3", "#3F37C9", "#0218F5", "#4361EE")## Blues

# Generate a ggplot
Sina_data %>% 
  filter(Dry_weight_ng > 1) %>%   # Filter the data for Dry_weight_ng greater than 1
  
  ggplot(aes(x = log(Dry_weight_µg), 
             y = log(CA_per_copepod_pmol))) +  # Define x and y aesthetics for the plot
  
  # Add points to the plot
  geom_point(
    aes(colour = Site),  # Color points by Site
    alpha = 0.2,         # Set point transparency
    size = 2,            # Set point size
    show.legend = TRUE  # Include points in the legend
  )+
  
  # Add a smooth line (linear regression model)
  geom_smooth(
    method = "lm",       # Fit a linear model
    se = TRUE,           # Show standard error
    alpha = 0.4,         # Set line transparency
    linewidth = 1,       # Set line width
    linetype = "solid",  # Set line type to solid
    show.legend = FALSE  # Exclude this from the legend
  )+
  
  # Set custom colors for the points
  scale_colour_manual(values = colours)+
  
  # Apply a custom theme (from the 'pubr' package)
  theme_pubr()+
  
  # Label the x-axis
  xlab(bquote(Ln~Dry~mass~(µg)))+
  
  # Label the y-axis
  ylab(bquote(Ln~Copepodamides~(pmol~ind^-1)))
```

###### Plot of the logarithmic relationship

```
# Attach the Sina_data dataframe (Note: Be cautious with attach() as it can cause naming conflicts)
attach(Sina_data)

# Define a custom function 'myfun' based on 'Dry_weight_µg'
myfun <- function(Dry_weight_µg) {
  (Dry_weight_µg^0.29604) + exp(0.25)
}

# Plot the curve generated by 'myfun' function
curve(myfun(x), from = 0, to = max(Dry_weight_µg))
```

```
# Define a vector of custom colors
colours <- c("#005F07","#00B60D", "#00FD12",
             "#86AD6C", "#008080", "#99E56A", 
             "#480CA8", "#3A0CA3", "#3F37C9", "#0218F5", "#4361EE")## Blues

# Generate a ggplot
Sina_data %>% 
  
  ggplot(aes(x = (Dry_weight_µg), 
             y = (CA_per_copepod_pmol))) +  # Define x and y aesthetics for the plot
  
  # Set y-axis limits, breaks, and expansion
  scale_y_continuous(limits = c(0,75),
                     breaks = c(0,15,30,45,60,75),
                     expand = c(0,0))+
  
  # Set x-axis limits, breaks, and expansion
  scale_x_continuous(limits = c(0, 1550),
                     breaks = c(0,250,500, 750, 1000, 1250, 1500),
                     expand = c(0.005,0))+
  
  # Add points to the plot
  geom_point(
    aes(colour = Site),  # Color points by Site
    alpha = 0.1,         # Set point transparency
    size = 1.5,          # Set point size
    show.legend = TRUE  # Include points in the legend
  )+
  
  # Add the function curve
  stat_function(fun = myfun, geom = "line")  +
  
  # Apply a custom theme (from the 'pubr' package)
  theme_pubr()+
  
  # Customize the legend position
  theme(legend.position = "right")+
  
  # Set custom colors for the legend
  scale_colour_manual(values = colours)+
  
  # Label the x-axis
  xlab(bquote(Dry~mass~(µg)))+
  
  # Label the y-axis
  ylab(bquote(Copepodamides~(pmol~ind^-1)))
```

###### Zoomed in

```
# Define a vector of custom colors
colours <- c("#005F07","#00B60D", "#00FD12",
             "#86AD6C", "#008080", "#99E56A",
             "#480CA8", "#3A0CA3", "#3F37C9", "#0218F5", "#4361EE")## Blues

# Generate a ggplot
Sina_data %>% 
  
  ggplot(aes(x = (Dry_weight_µg), 
             y = (CA_per_copepod_pmol))) +  # Define x and y aesthetics for the plot
  
  # Set y-axis limits, breaks, and expansion
  scale_y_continuous(limits = c(0,75),
                     breaks = c(0,5,10,15,20),
                     expand = c(0,0))+
  
  # Set x-axis limits, breaks, and expansion
  scale_x_continuous(limits = c(0, 755),
                     breaks = c(0,250,500, 750),
                     expand = c(0.005,0))+
  
  # Adjust the y-axis limits
  coord_cartesian(ylim = c(0,20))+
  
  # Add points to the plot
  geom_point(
    aes(colour = Site),  # Color points by Site
    alpha = 0.2,         # Set point transparency
    size = 2,            # Set point size
    show.legend = TRUE  # Include points in the legend
  )+
  
  # Set custom colors for the legend
  scale_colour_manual(values = colours)+
  
  # Add the function curve
  stat_function(fun = myfun, geom = "line")  +
  
  # Apply a custom theme (from the 'pubr' package)
  theme_pubr()+
  
  # Label the x-axis
  xlab(bquote(Dry~mass~(µg)))+
  
  # Label the y-axis
  ylab(bquote(Copepodamides~(pmol~ind^-1)))
```

###### More zoomed in

```
# Define a vector of custom colors
colours <- c("#005F07","#00B60D", "#00FD12",
             "#86AD6C", "#008080", "#99E56A",
             "#480CA8", "#3A0CA3", "#3F37C9", "#0218F5", "#4361EE")## Blues

# Generate a ggplot
Sina_data %>% 
  
  ggplot(aes(x = (Dry_weight_µg), 
             y = (CA_per_copepod_pmol))) +  # Define x and y aesthetics for the plot
  
  # Set y-axis limits, breaks, and expansion
  scale_y_continuous(limits = c(0,75),
                     breaks = c(0,5,10,15,20,25,30),
                     expand = c(0,0))+
  
  # Set x-axis limits, breaks, and expansion
  scale_x_continuous(limits = c(0, 80),
                     breaks = c(0,20,40,60,80),
                     expand = c(0.005,0))+
  
  # Adjust the y-axis limits
  coord_cartesian(ylim = c(0,30))+
  
  # Add points to the plot
  geom_point(
    aes(colour = Site),  # Color points by Site
    alpha = 0.2,         # Set point transparency
    size = 2,            # Set point size
    show.legend = TRUE  # Include points in the legend
  )+
  
  # Set custom colors for the legend
  scale_colour_manual(values = colours)+
  
  # Add the function curve
  stat_function(fun = myfun, geom = "line")  +
  
  # Apply a custom theme (from the 'pubr' package)
  theme_pubr()+
  
  # Label the x-axis
  xlab(bquote(Dry~mass~(µg)))+
  
  # Label the y-axis
  ylab(bquote(Copepodamides~(pmol~ind^-1)))
```

###### Results ANCOVA

Both the statistical analyses (ANCOVA) and the visualizations clearly
suggests that copepod size/mass is the main determining factor of
copepodamide content/amount for both marine and freshwater copepods.
Even after controlling for their size, habitat type does not
significantly modulate this relationship.

#### Assessing the effect of time between sampling and extraction on total copepodamide content.

##### The model

```
# Fit a linear model using Time_diff_min as predictor
model_time <- lm(log(CA_per_copepod_pmol) ~ Time_diff_min, data = Sina_data)

# Fit a linear model using Time_diff_min and Dry_weight_µg as predictors
model_time_weight <- lm(log(CA_per_copepod_pmol) ~ Time_diff_min + Dry_weight_µg, data = Sina_data)

# Summarize the first model
summary.lm(model_time)
```

```
## 
## Call:
## lm(formula = log(CA_per_copepod_pmol) ~ Time_diff_min, data = Sina_data)
## 
## Residuals:
##     Min      1Q  Median      3Q     Max 
## -5.5224 -1.0008  0.2905  1.1100  3.1350 
## 
## Coefficients:
##                Estimate Std. Error t value Pr(>|t|)    
## (Intercept)    1.329442   0.294572   4.513 1.56e-05 ***
## Time_diff_min -0.001963   0.001529  -1.284    0.202    
## ---
## Signif. codes:  0 '***' 0.001 '**' 0.01 '*' 0.05 '.' 0.1 ' ' 1
## 
## Residual standard error: 1.572 on 114 degrees of freedom
## Multiple R-squared:  0.01426,    Adjusted R-squared:  0.005612 
## F-statistic: 1.649 on 1 and 114 DF,  p-value: 0.2017
```

```
# Summarize the secondmodel
summary.lm(model_time_weight)
```

```
## 
## Call:
## lm(formula = log(CA_per_copepod_pmol) ~ Time_diff_min + Dry_weight_µg, 
##     data = Sina_data)
## 
## Residuals:
##    Min     1Q Median     3Q    Max 
## -5.346 -1.041  0.312  1.103  2.812 
## 
## Coefficients:
##                Estimate Std. Error t value Pr(>|t|)    
## (Intercept)    1.337806   0.286007   4.678  8.1e-06 ***
## Time_diff_min -0.002860   0.001518  -1.884   0.0621 .  
## Dry_weight_µg  0.002570   0.000912   2.818   0.0057 ** 
## ---
## Signif. codes:  0 '***' 0.001 '**' 0.01 '*' 0.05 '.' 0.1 ' ' 1
## 
## Residual standard error: 1.526 on 113 degrees of freedom
## Multiple R-squared:  0.079,  Adjusted R-squared:  0.06269 
## F-statistic: 4.846 on 2 and 113 DF,  p-value: 0.009567
```

###### Table of results

```
# Create a regression table for the model_time
t1 <- tbl_regression(model_time,
                     intercept = TRUE, 
                     estimate_fun = ~style_number(.x, digits = 4),
                     label = list("(Intercept)" ~ "Intercept",
                                  Time_diff_min ~ "Time (min)")) %>% 
      modify_header(label = "**Variable**") %>% 
      bold_labels() %>% 
      bold_p()  %>% 
      add_glance_table(include = c(r.squared, df, df.residual))

# Create a regression table for the model_time_weight
t2 <- tbl_regression(model_time_weight,
                     intercept = TRUE, 
                     estimate_fun = ~style_number(.x, digits = 4),
                     label = list("(Intercept)" ~ "Intercept",
                                  Time_diff_min ~ "Time (min)",
                                  Dry_weight_µg ~ "Dry mass (µg)")) %>% 
      modify_header(label = "**Variable**") %>% 
      bold_labels() %>% 
      bold_p()%>% 
      add_glance_table(include = c(r.squared, df, df.residual))

# Merge the two regression tables with appropriate headers
merged_table <- tbl_merge(
                  tbls = list(t2, t1),
                  tab_spanner = c("**Corrected for copepod mass**", "**Raw time effect**"))

# Display the merged table
merged_table
```

| **Variable** | **Corrected for copepod mass** | | | **Raw time effect** | | |
| --- | --- | --- | --- | --- | --- | --- |
|  | **Beta** | **95% CI**^1^ | **p-value** | **Beta** | **95% CI**^1^ | **p-value** |
| Intercept | 1.3378 | 0.7712, 1.9044 | <0.001 | 1.3294 | 0.7459, 1.9130 | <0.001 |
| Time (min) | -0.0029 | -0.0059, 0.0001 | 0.062 | -0.0020 | -0.0050, 0.0011 | 0.2 |
| Dry mass (µg) | 0.0026 | 0.0008, 0.0044 | 0.006 |  |  |  |
| R² | 0.079 |  |  | 0.014 |  |  |
| df | 2 |  |  | 1 |  |  |
| Residual df | 113 |  |  | 114 |  |  |
|  |  |  |  |  |  |  |
| --- | --- | --- | --- | --- | --- | --- |
| ^1^ CI = Confidence Interval | | | | | | |

```
# Save the table as an image

## merged_table %>% as_gt() %>% gt::gtsave("table_time_on_CA.png")
```

Results: Non-significant negative trend of time between sampling
& extraction on CA content, regardless of if dry mass is accounted
for or not.

##### Plot of CA content as a function of time between sampling and extraction (min), points scaled by dry mass (µg)

```
# Define a vector of color codes
colours <- c("#005F07","#00B60D", "#00FD12",
             "#86AD6C", "#008080", "#99E56A",  ##Greens
             "#480CA8", "#3F37C9", "#0218F5", "#4361EE") ## Blues

# Create a ggplot object using Sina_data, filtering by Dry_weight_ng > 1
Sina_data %>% filter(Dry_weight_ng > 1) %>% 
  ggplot(aes(x = (Time_diff_min), 
             y = log(CA_per_copepod_pmol))) +
  
  # Add points to the plot, adjusting size and color by Dry_weight_µg, and color by Site
  geom_point(aes(size = Dry_weight_µg,
                 colour = Site),
             alpha = 0.3,
             show.legend = TRUE)+
  
  # Add a smooth regression line to the plot
  geom_smooth(colour = "black",
              method = "lm",
              se = TRUE,
              alpha = 0.4,
              linewidth = 1,
              linetype = "solid",
              show.legend = FALSE)+
  
  # Customize color scale
  scale_colour_manual(values = colours)+
  
  # Customize size scale
  scale_size_continuous(name = "Dry mass (µg)",
                        breaks = c(250,500, 750, 1000,1250))+
  
  # Apply the 'theme_pubr' theme with legend positioned on the right
  theme_pubr(legend =  "right")+
  
  # Customize legend appearance
  theme(legend.title = element_text(size = 10,
                                    colour = "black"),
        legend.text = element_text(size =10,
                                    colour = "black"),
        legend.spacing.x = unit(0.1, 'mm'),
        legend.spacing.y = unit(0.1, "mm"))+
  
  # Customize appearance of color legend symbols
  guides(colour = guide_legend(override.aes = list(size = 3.5)))+ 
  
  # Label the x-axis
  xlab(bquote(Time~between~sampling~and~extraction~(minutes)))+
  
  # Label the y-axis
  ylab(bquote(Ln~Copepodamides~(pmol~ind^-1)))
```

```
# Save the plot as an image named "CA_over_time_SizeScaled.png"
## ggsave("CA_over_time_SizeScaled.png", width = 170, height = 120, units = "mm", dpi=700)
```

### Non-targeted copepodamide analysis

#### Data import and preparation

```
# Read the data 
Nontargeted_CA <- read_delim("Arnoldt_untargeted_analysis_data.csv",
                            escape_double = FALSE, # Data is in percentages
                            trim_ws = TRUE)

# Attach the data for easier referencing of variables (Note: Using attach can lead to potential issues, consider using with() or other alternatives)
attach(Nontargeted_CA)

# Create a new dataframe data_id_nontarget containing columns Site and Type
data_id_nontarget <- Nontargeted_CA %>% 
  select(Site, Type)

# Create a new dataframe data_nMDS_nontarget containing all columns except Site and Type
data_nMDS_nontarget <- Nontargeted_CA %>% 
  select(!Site & !Type)

# Create a new dataframe data_nMDS_nontarget_props by dividing each element in data_nMDS_nontarget by the row sums
data_nMDS_nontarget_props <- data_nMDS_nontarget/rowSums(data_nMDS_nontarget)
```

#### Ordination (nMDS)

```
# Set the seed for reproducibility
set.seed(42)

# Perform non-metric multidimensional scaling (nMDS)
nMDS_nontargeted_props <- metaMDS(data_nMDS_nontarget_props,
                distance = "bray",  # Use Bray-Curtis dissimilarity
                k = 2,              # Number of dimensions for nMDS
                try = 99,           # Maximum number of random starts to try
                trymax = 200,       # Maximum number of iterations
                autotransform = FALSE)  # Disable automatic data transformation
```

```
# Show results of nMDS
nMDS_nontargeted_props
```

```
## 
## Call:
## metaMDS(comm = data_nMDS_nontarget_props, distance = "bray",      k = 2, try = 99, trymax = 200, autotransform = FALSE) 
## 
## global Multidimensional Scaling using monoMDS
## 
## Data:     data_nMDS_nontarget_props 
## Distance: bray 
## 
## Dimensions: 2 
## Stress:     0.04206032 
## Stress type 1, weak ties
## Best solution was repeated 17 times in 99 tries
## The best solution was from try 22 (random start)
## Scaling: centring, PC rotation, halfchange scaling 
## Species: expanded scores based on 'data_nMDS_nontarget_props'
```

#### Plotting the nMDS

```
# Extract the site scores from 'nMDS_nontargeted_props' and convert to a data frame
datascores_nontarget_props <- as.data.frame(scores(nMDS_nontargeted_props, display = "sites"))

# Combine the data frame 'data_id_nontarget' with 'datascores_nontarget_props' using column binding
scores_nontarget_props <- cbind(data_id_nontarget, datascores_nontarget_props)

# Calculate the centroids (means) of 'NMDS1' and 'NMDS2' for each unique value of 'Type' in 'scores_nontarget_props'
centroids_type_nontarget_props <- aggregate(cbind(NMDS1, NMDS2) ~ Type, data = scores_nontarget_props, FUN = mean)

# Define a vector of color codes used for plotting
colours <- c("#005F07","#00B60D", "#00FD12",
             "#86AD6C", "#008080", "#99E56A", "#228B22", 
             "#0E1DFF", "#2472FA", "#5713FF", "#657BC5", "#0000D1")  # Blues/purples

# Initialize a ggplot object with 'NMDS1' on the x-axis and 'NMDS2' on the y-axis
scores_nontarget_props %>% 
  ggplot(aes(x = NMDS1, y = NMDS2)) + 
  
  # Add confidence ellipses to the plot
  stat_ellipse(aes(fill = Type),  # Color fill based on Type
               geom = "polygon",  # Draw ellipses as polygons
               alpha = 0.07,  # Transparency level
               linewidth = 0.5,  # Line width of the ellipse
               level = 0.95,  # Confidence level
               show.legend = FALSE) +  # Do not show legend for this layer
  
  # Add points to the plot
  geom_point(aes(fill = Site),  # Color fill based on Site
             colour = "black",  # Color of point outline
             size = 2,  # Size of points
             shape = 21,  # Shape of points (filled circle)
             alpha = 0.6,  # Transparency level
             show.legend = FALSE) +  # Do not show legend for this layer
  
  # Add points representing centroids
  geom_point(aes(fill = Type),  # Color fill based on Type
             data = centroids_type_nontarget_props,  # Data source for centroids
             size = 4,  # Size of points representing centroids
             shape = 21,  # Shape of points (filled circle)
             alpha = 0.9,  # Transparency level
             colour = "black",  # Color of point outline
             show.legend = TRUE) +  # Show legend for this layer

# Apply a classic theme to the plot
theme_classic() +

# Set properties for the x-axis
scale_x_continuous(limits = c(-1.7, 1.7),  # Set x-axis limits
                   expand = c(0,0),  # Set axis expansion
                   breaks = c(-1.7, -0.8, 0, 0.8, 1.7)) +  # Set breaks for x-axis

# Set manual fill colors and legend title for 'Site' variable
scale_fill_manual(values = colours, name = "Site", 
                  labels = c("F1", "F2", "F3", "F4", "F5", "F6", "Fre", 
                             "M1", "M2", "M3", "M4", "Mar")) +

# Set manual color scale for the legend
scale_colour_manual(values = c("#228B22", "#0000D1")) +

# Remove color legend
guides(colour = "none") +

# Customize various aspects of the plot's appearance
theme(
  plot.margin = unit(c(0.3, 0.2, 0.1, 0.1), "cm"),  # Set plot margins
  axis.title.y = element_text(vjust = 0, size = 8),  # Customize y-axis title
  axis.title.x = element_text(size = 8),  # Customize x-axis title
  axis.text.y = element_text(colour = "black", size = 8),  # Customize y-axis text
  axis.text.x = element_text(colour = "black", size = 8),  # Customize x-axis text
  legend.text = element_text(colour = "black", size = 7),  # Customize legend text
  legend.title = element_text(colour = "black", size = 8),  # Customize legend title
  legend.position = c(0.08,0.38),  # Set legend position
  legend.margin = margin(unit(c(0, -23, 0, -25), "cm")),  # Set legend margins
  legend.key.size = unit(0.3, 'cm')  # Set size of legend keys
) +

# Set properties for the fill legend
guides(fill = guide_legend(override.aes = list(size = 2.5, alpha = 0.7)))
```

```
# Save the plot with specified dimensions and resolution
## ggsave("Non-targeted_nMDS_new_props.tiff", width = 80, height = 65, units = "mm", dpi=700)
```

#### Multivariate analysis - PERMANOVA

```
# Set seed for reproducibility
set.seed(42)

# Testing for homogeneity of dispersion

# Calculate Bray-Curtis dissimilarity matrix for nontargeted data
nontargeted_dist_props <- vegdist(data_nMDS_nontarget_props, method = "bray")

# Perform multivariate homogeneity of group dispersions test
dispersion_nontargeted_props <- betadisper(nontargeted_dist_props, group = data_id_nontarget$Type)

# Display results of the betadisper test
dispersion_nontargeted_props
```

```
## 
##  Homogeneity of multivariate dispersions
## 
## Call: betadisper(d = nontargeted_dist_props, group =
## data_id_nontarget$Type)
## 
## No. of Positive Eigenvalues: 8
## No. of Negative Eigenvalues: 1
## 
## Average distance to median:
## Freshwater     Marine 
##     0.2175     0.2609 
## 
## Eigenvalues for PCoA axes:
## (Showing 8 of 9 eigenvalues)
##    PCoA1    PCoA2    PCoA3    PCoA4    PCoA5    PCoA6    PCoA7    PCoA8 
## 0.805510 0.167855 0.151856 0.104759 0.055685 0.029860 0.009802 0.001861
```

```
# Plot the results of the betadisper test
plot(dispersion_nontargeted_props)
```

```
# Perform ANOVA on the betadisper results
anova(dispersion_nontargeted_props)
```

```
## # A tibble: 2 × 5
##      Df `Sum Sq` `Mean Sq` `F value` `Pr(>F)`
##   <int>    <dbl>     <dbl>     <dbl>    <dbl>
## 1     1  0.00453   0.00453     0.552    0.479
## 2     8  0.0656    0.00820    NA       NA
```

```
# Perform permutation test on the betadisper results
permutest(dispersion_nontargeted_props, permutations = 9999)
```

```
## 
## Permutation test for homogeneity of multivariate dispersions
## Permutation: free
## Number of permutations: 9999
## 
## Response: Distances
##           Df   Sum Sq   Mean Sq      F N.Perm Pr(>F)
## Groups     1 0.004527 0.0045270 0.5524   9999 0.4658
## Residuals  8 0.065567 0.0081959
```

```
# Conduct Tukey's Honestly Significant Difference (HSD) test on the betadisper results
TukeyHSD(dispersion_nontargeted_props)
```

```
##   Tukey multiple comparisons of means
##     95% family-wise confidence level
## 
## Fit: aov(formula = distances ~ group, data = df)
## 
## $group
##                         diff         lwr       upr     p adj
## Marine-Freshwater 0.04343094 -0.09132618 0.1781881 0.4786133
```

```
# Bootstrapping and testing for differences between the groups

# Fit a PERMANOVA model using Bray-Curtis dissimilarities
fit_nontarget_props <- adonis2(data_nMDS_nontarget_props ~ data_id_nontarget$Type, 
                               data=data_id_nontarget, 
                               permutations=9999, 
                               method="bray")

# Display results of the PERMANOVA test
fit_nontarget_props
```

```
## # A tibble: 3 × 5
##      Df SumOfSqs    R2     F `Pr(>F)`
##   <dbl>    <dbl> <dbl> <dbl>    <dbl>
## 1     1    0.718 0.545  9.57   0.0038
## 2     8    0.600 0.455 NA     NA     
## 3     9    1.32  1     NA     NA
```

```
# Calculate Omega Squared (partial R-squared) for the PERMANOVA model
adonis_OmegaSq(fit_nontarget_props, partial = TRUE)
```

```
## # A tibble: 3 × 5
##      Df SumOfSqs     F parOmegaSq `Pr(>F)`
##   <dbl>    <dbl> <dbl>      <dbl>    <dbl>
## 1     1    0.718  9.57      0.462   0.0038
## 2     8    0.600 NA        NA      NA     
## 3     9    1.32  NA        NA      NA
```

```
# Simper analysis for nontargeted data
simper_nontargeted_props <- simper(data_nMDS_nontarget_props, data_id_nontarget$Type)

# Display summary of simper results, ordered by importance
summary(simper_nontargeted_props, ordered = TRUE)
```

```
## 
## Contrast: Freshwater_Marine 
## 
##            average      sd   ratio     ava     avb cumsum     p    
## 686.5_dhCA 0.06318 0.04206 1.50200 0.16123 0.03562  0.099 0.014 *  
## 712.6_dhCA 0.05337 0.02603 2.05000 0.11849 0.01175  0.182 0.007 ** 
## 714.6_dhCA 0.04816 0.01770 2.72100 0.11047 0.01415  0.257 0.006 ** 
## 708.6_dhCA 0.04806 0.03176 1.51300 0.06423 0.15615  0.332 0.106    
## 706.6_CA   0.04346 0.03121 1.39300 0.00000 0.08693  0.400 0.001 ***
## 710.6_dhCA 0.03868 0.02196 1.76100 0.12143 0.04707  0.460 0.026 *  
## 732.6_CA   0.03592 0.03446 1.04200 0.00000 0.07183  0.517 0.001 ***
## 688.6_dhCA 0.03207 0.03042 1.05400 0.08818 0.07684  0.567 0.462    
## 708.6_CA   0.03118 0.02771 1.12500 0.00000 0.06237  0.615 0.001 ***
## 760.6_dhCA 0.03094 0.01977 1.56500 0.05229 0.06879  0.664 0.491    
## 660.5_dhCA 0.02638 0.01819 1.45000 0.08170 0.04526  0.705 0.214    
## 706.5_dhCA 0.02490 0.00761 3.27000 0.00000 0.04980  0.744 0.001 ***
## 758.7_CA   0.02066 0.01796 1.15000 0.00000 0.04132  0.776 0.001 ***
## 734.6_dhCA 0.01831 0.01560 1.17400 0.10574 0.07307  0.805 0.327    
## 674.5_dhCA 0.01494 0.00853 1.75100 0.03135 0.00219  0.828 0.020 *  
## 736.6_dhCA 0.01354 0.00752 1.80100 0.02708 0.00000  0.849 0.019 *  
## 760.7_CA   0.01107 0.01959 0.56500 0.00000 0.02215  0.866 0.001 ***
## 734.6_CA   0.01027 0.01243 0.82600 0.00000 0.02054  0.882 0.001 ***
## 686.6_CA   0.00936 0.00603 1.55200 0.00000 0.01871  0.897 0.001 ***
## 710.6_CA   0.00894 0.01151 0.77700 0.00000 0.01789  0.911 0.001 ***
## 680.5_dhCA 0.00692 0.00716 0.96700 0.00000 0.01384  0.922 0.001 ***
## 688.6_CA   0.00654 0.01158 0.56500 0.00000 0.01309  0.932 0.001 ***
## 700.5_dhCA 0.00523 0.00768 0.68100 0.01046 0.00000  0.940 0.381    
## 704.5_CA   0.00487 0.00433 1.12400 0.00000 0.00973  0.948 0.001 ***
## 702.7_dhCA 0.00444 0.00478 0.92900 0.00888 0.00000  0.955 0.231    
## 712.6_CA   0.00361 0.00216 1.67300 0.00000 0.00721  0.960 0.001 ***
## 684.5_dhCA 0.00357 0.00407 0.87800 0.00641 0.00219  0.966 0.676    
## 660.5_CA   0.00325 0.00574 0.56500 0.00000 0.00649  0.971 0.001 ***
## 684.4_CA   0.00300 0.00188 1.59200 0.00000 0.00599  0.976 0.001 ***
## 658.5_CA   0.00279 0.00375 0.74500 0.00000 0.00559  0.980 0.001 ***
## 682.5_dhCA 0.00236 0.00260 0.90600 0.00000 0.00472  0.984 0.001 ***
## 738.7_dhCA 0.00217 0.00358 0.60700 0.00435 0.00000  0.987 0.381    
## 762.6_dhCA 0.00172 0.00289 0.59400 0.00344 0.00000  0.990 0.406    
## 678.5_CA   0.00140 0.00247 0.56500 0.00000 0.00279  0.992 0.001 ***
## 672.5_dhCA 0.00134 0.00307 0.43800 0.00269 0.00000  0.994 0.381    
## 714.6_CA   0.00127 0.00224 0.56500 0.00000 0.00253  0.996 0.001 ***
## 742.6_dhCA 0.00086 0.00152 0.56500 0.00000 0.00172  0.998 0.001 ***
## 716.5_CA   0.00084 0.00148 0.56500 0.00000 0.00167  0.999 0.001 ***
## 702.4_dhCA 0.00080 0.00182 0.43800 0.00159 0.00000  1.000 0.406    
## ---
## Signif. codes:  0 '***' 0.001 '**' 0.01 '*' 0.05 '.' 0.1 ' ' 1
## Permutation: free
## Number of permutations: 999
```

### Stacked barchart of copepodamide composition

#### Data wrangling

```
# Create a new data frame 'data_id_long' by selecting specific columns from 'data_id_target'
data_id_long <- data_id_target %>% select(Sample, Site, Type, Taxon)

# Combine 'data_id_long' with 'data_nMDS_target_props' using column binding
data_stacked <- cbind(data_id_long, data_nMDS_target_props)

# Perform a pivot_longer operation on the data, creating a new column 'Copepodamide_species'
Sina_long <- data_stacked %>% 
  pivot_longer(!Sample:Taxon, names_to = "Copepodamide_species") %>% 
  as.data.frame() 

# Convert 'Copepodamide_species' column to a factor with specific levels and labels
Sina_long$Copepodamide_species <- factor(Sina_long$Copepodamide_species,
                                         levels = c("448.3_CA", 
                                                    "686.5_CA", 
                                                    "706.6_CA", 
                                                    "708.6_CA",
                                                    "712.5_CA", 
                                                    "732.5_CA", 
                                                    "734.5_CA", 
                                                    "758.6_CA", 
                                                    "450.3_dhCA",
                                                    "660.5_dhCA", 
                                                    "674.6_dhCA",
                                                    "680.5_dhCA", 
                                                    "686.5_dhCA",
                                                    "688.6_dhCA", 
                                                    "706.5_dhCA", 
                                                    "710.6_dhCA", 
                                                    "708.5_dhCA",
                                                    "712.6_dhCA",
                                                    "714.6_dhCA", 
                                                    "734.5_dhCA", 
                                                    "736.6_dhCA", 
                                                    "760.5_dhCA"),
                                         
                                         labels = c("lyso-CA", 
                                                    "16:0-CA", 
                                                    "18:4-CA", 
                                                    "18:3-CA", 
                                                    "18:1-CA", 
                                                    "20:5-CA", 
                                                    "20:4-CA", 
                                                    "22:6-CA", 
                                                    "lyso-dhCA",
                                                    "14:0-dhCA", 
                                                    "15:0-dhCA", 
                                                    "16:4-dhCA", 
                                                    "16:1-dhCA", 
                                                    "16:0-dhCA", 
                                                    "18:5-dhCA", 
                                                    "18:4-dhCA",
                                                    "18:3-dhCA",
                                                    "18:2-dhCA",
                                                    "18:1-dhCA",
                                                    "20:5-dhCA",
                                                    "20:4-dhCA",
                                                    "22:6-dhCA"))

# Display the structure of the 'Sina_long' data frame
str(Sina_long)
```

```
## 'data.frame':    2552 obs. of  6 variables:
##  $ Sample              : chr  "F1a" "F1a" "F1a" "F1a" ...
##  $ Site                : chr  "F1" "F1" "F1" "F1" ...
##  $ Type                : chr  "Freshwater" "Freshwater" "Freshwater" "Freshwater" ...
##  $ Taxon               : chr  "Cyclops strenuus" "Cyclops strenuus" "Cyclops strenuus" "Cyclops strenuus" ...
##  $ Copepodamide_species: Factor w/ 22 levels "lyso-CA","16:0-CA",..: 1 9 10 11 12 13 2 14 15 3 ...
##  $ value               : num  0 0.0207 0.0486 0.0326 0 ...
```

```
# Display the first few rows of 'Sina_long'
head(Sina_long)
```

```
## # A tibble: 6 × 6
##   Sample Site  Type       Taxon            Copepodamide_species  value
##   <chr>  <chr> <chr>      <chr>            <fct>                 <dbl>
## 1 F1a    F1    Freshwater Cyclops strenuus lyso-CA              0     
## 2 F1a    F1    Freshwater Cyclops strenuus lyso-dhCA            0.0207
## 3 F1a    F1    Freshwater Cyclops strenuus 14:0-dhCA            0.0486
## 4 F1a    F1    Freshwater Cyclops strenuus 15:0-dhCA            0.0326
## 5 F1a    F1    Freshwater Cyclops strenuus 16:4-dhCA            0     
## 6 F1a    F1    Freshwater Cyclops strenuus 16:1-dhCA            0.456
```

#### Grouped by type & taxa

```
# Define a vector of color codes used for plotting
colours <- c("#2999BA", "#0990BA", "#0980BA", 
             "#0970BA", "#0960BA", "#0950BA", 
             "#0920BA", "#0900BA", "#B2E972", 
             "#A7E06C", "#A7E04C","#9CD866", 
             "#5CD810", "#158902", "#157502", 
             "#156912", "#156212", "#155812", 
             "#155112", "#154712", "#153912", "#153012")

# Create a ggplot object with specific aesthetics and data
Sina_long  %>% 
  ggplot(aes(y = interaction(Sample, Taxon, Type),  # Define y-axis as interaction of Sample, Taxon, and Type
             x = value,  # Define x-axis as 'value'
             fill = Copepodamide_species)) +  # Fill colors based on 'Copepodamide_species'
  
  # Add bar geometry to the plot
  geom_bar(position = position_fill(reverse = TRUE),  # Position bars using fill, reverse order
           stat = "identity",  # Use actual values as heights
           width = 1,  # Set width of bars
           show.legend = TRUE) +  # Show legend for this layer
  
  # Set properties for the x-axis
  scale_x_continuous(limits = c(0,1),  # Set x-axis limits
                     labels = c(0, 25, 50, 75, 100),  # Set x-axis labels
                     expand = c(0,0)) +  # Set axis expansion
  
  # Set properties for the y-axis
  scale_y_discrete(guide = "axis_nested",  # Use nested axis guide for y-axis
                   name = "Habitat") +  # Set y-axis label
  
  xlab("Relative abundance (%)") +  # Set x-axis label
  
  # Set manual fill colors and legend title for 'Copepodamide_species'
  scale_fill_manual(values = colours,
                    name = "Copepodamide") +
  
  # Apply a classic theme to the plot
  theme_classic()  +
  
  # Customize various aspects of the plot's appearance
  theme(
    legend.position = "right",  # Set legend position
    legend.title = element_text(size = 9),  # Set legend title size
    legend.text = element_text(size = 9),  # Set legend text size
    legend.direction = "vertical",  # Set legend direction
    text = element_text(colour = "black"),  # Set text color
    axis.title.y = element_blank(),  # Remove y-axis title
    axis.title.x = element_text(size = 8),  # Set x-axis title size
    axis.text.y  = element_text(colour = "black",  # Set y-axis text color, size, and face
                                  size = 7,
                                  face = "plain"),
    axis.text.x  = element_text(colour = "black",  # Set x-axis text color and size
                                  size = 8),
    plot.margin = unit(c(0.2, 0.2, 0.1, 0.1), "cm"),  # Set plot margins
    legend.margin = margin(unit(c(0, 1, 0, -2), "cm")),  # Set legend margins
    legend.key.size = unit(0.5, 'cm')  # Set size of legend keys
  ) +
  
  guides(fill=guide_legend(ncol=1))  # Set properties for the fill legend
```

```
# Save the plot with specified dimensions and resolution

## ggsave("Copepodamide_composition_taxa.png", width = 130, height = 235, units = "mm", dpi=700)
```

#### Grouped by type and site

```
# Define a vector of color codes used for plotting
colours <- c("#2999BA", "#0990BA", "#0980BA", 
             "#0970BA", "#0960BA", "#0950BA",
             "#0920BA", "#0900BA", "#B2E972",
             "#A7E06C", "#A7E04C","#9CD866",
             "#5CD810", "#158902", "#157502", 
             "#156912", "#156212", "#155812", 
             "#155112", "#154712", "#153912", "#153012")

# Create a ggplot object with specific aesthetics and data
Sina_long  %>% 
  ggplot(aes(y = interaction(Sample, Site, Type),  # Define y-axis as interaction of Sample, Site, and Type
             x = value,  # Define x-axis as 'value'
             fill = Copepodamide_species)) +  # Fill colors based on 'Copepodamide_species'
  
  # Add bar geometry to the plot
  geom_bar(position = position_fill(reverse = TRUE),  # Position bars using fill, reverse order
           stat = "identity",  # Use actual values as heights
           width = 1,  # Set width of bars
           show.legend = TRUE) +  # Show legend for this layer
  
  # Set properties for the x-axis
  scale_x_continuous(limits = c(0,1),  # Set x-axis limits
                     labels = c(0, 25, 50, 75, 100),  # Set x-axis labels
                     expand = c(0,0)) +  # Set axis expansion
  
  # Set properties for the y-axis
  scale_y_discrete(guide = "axis_nested",  # Use nested axis guide for y-axis
                   name = "Habitat") +  # Set y-axis label
  
  xlab("Relative abundance (%)") +  # Set x-axis label
  
  # Set manual fill colors and legend title for 'Copepodamide_species'
  scale_fill_manual(values = colours,
                    name = "Copepodamide") +
  
  # Apply a classic theme to the plot
  theme_classic()  +
  
  # Customize various aspects of the plot's appearance
  theme(
    legend.position = "right",  # Set legend position
    legend.title = element_text(size = 8),  # Set legend title size
    legend.text = element_text(size = 8),  # Set legend text size
    legend.direction = "vertical",  # Set legend direction
    text = element_text(colour = "black"),  # Set text color
    axis.title.y = element_blank(),  # Remove y-axis title
    axis.title.x = element_text(size = 8),  # Set x-axis title size
    axis.text.y  = element_text(colour = "black",  # Set y-axis text color, size, and face
                                  size = 7,
                                  face = "plain"),
    axis.text.x  = element_text(colour = "black",  # Set x-axis text color and size
                                  size = 8),
    plot.margin = unit(c(0.2, 0.2, 0.1, 0.1), "cm"),  # Set plot margins
    legend.margin = margin(unit(c(0, 1, 0, -2), "cm")),  # Set legend margins
    legend.key.size = unit(0.5, 'cm')  # Set size of legend keys
  ) +
  
  guides(fill=guide_legend(ncol=1))  # Set properties for the fill legend
```

```
# Save the plot as a TIFF file with specified dimensions and resolution
## ggsave("Copepodamide_composition_site.tiff", width = 100, height = 240, units = "mm", dpi=700)
```

### Additional plots

#### Counts of taxa by habitat

##### Create dataframe

```
# Calculate the count of occurrences for each combination of 'Type' and 'Taxon'
# Calculate the percentage of each combination relative to the total count
taxon_counts <-
  Sina_data %>% 
  select(Type, Taxon) %>%  # Select 'Type' and 'Taxon' columns
  group_by(Type, Taxon) %>%  # Group the data by 'Type' and 'Taxon'
  summarize(Count = n()) %>%  # Calculate the count of occurrences
  mutate(Percentage = round(x = (100*(Count / sum(Count))), digits = 1))  # Calculate the percentage
```

##### Plot for freshwater samples

```
# Filter 'taxon_counts' data for 'Type' equal to "Freshwater"
# Reorder 'Taxon' based on descending 'Percentage' values
# Convert the result to a data frame
taxon_counts %>% 
  filter(Type == "Freshwater") %>%  
  mutate(Taxon = reorder(Taxon, -Percentage)) %>%  
  data.frame() %>%  

ggplot(aes(x = Taxon, 
           y = Percentage)) +  

  # Set properties for the y-axis
  scale_y_continuous(
    expand = c(0,0),  # Set expansion of y-axis
    limits = c(0,52),  # Set limits of y-axis
    breaks = c(0, 5,10,15,20,25,30,35,40,45,50)  # Set breaks for y-axis
  )+  

  # Set properties for the x-axis labels
  scale_x_discrete(
    labels = c(
      expression(italic("Cyclops strenuus")),
      expression(italic("Cyclops sp.")), 
      expression("Unknown"),
      expression(italic("Eudiaptomus graciloides")),
      expression("Cyclopoid copepod"), 
      expression(italic("Macrocyclops albidus"))
    )
  )+
  
  # Add a bar plot layer using actual values as heights, with specified colors
  geom_bar(
    stat = "identity",  # Use actual values for heights
    fill = "grey60",  # Set fill color of bars
    colour = "grey40"  # Set border color of bars
  )+  

  # Add labels for the title, x-axis, and y-axis
  labs(
    title = "Freshwater, n = 60",  # Set plot title
    x = "",  # Set x-axis label to empty
    y = "Proportion (%)"  # Set y-axis label
  )+
  
  # Add text labels to the bar plot
  geom_text(
    aes(label = Count),  # Use 'Count' for labels
    vjust = -0.4,  # Adjust vertical position of labels
    size = 4,  # Set text size
    color = "black"  # Set text color
  ) +  
  
  # Apply a classic theme to the plot
  theme_classic()+  

  # Customize various aspects of the plot's appearance
  theme(
    axis.text = element_text(size = 9, colour = "black"),  # Customize axis text
    axis.text.x = element_text(size = 9, angle = 50, hjust = 1),  # Customize x-axis text
    axis.title.y = element_text(size = 12, vjust = 1.8, colour = "black")  # Customize y-axis title
  )
```

```
# Save the plot as a PNG file with specified dimensions and resolution
## ggsave("Taxa_comp_FW.png", width = 100, height = 100, units = "mm", dpi=700)
```

##### Plot for marine samples

```
# Filter the 'taxon_counts' data to include only entries with Type "Marine"
# Reorder 'Taxon' based on descending 'Percentage' values
# Convert the result to a data frame
taxon_counts %>% 
  filter(Type == "Marine") %>%  
  mutate(Taxon = reorder(Taxon, -Percentage)) %>%  
  data.frame() %>%  

# Create a ggplot object with specified aesthetics
ggplot(aes(x = Taxon, 
           y = Percentage)) +  

  # Customize the y-axis scale properties
  scale_y_continuous(
    expand = c(0,0),  # Set expansion of y-axis
    limits = c(0,40),  # Set limits of y-axis
    breaks = c(0, 5,10,15,20,25,30,35,40)  # Set breaks for y-axis
  )+  

  # Customize the x-axis labels
  scale_x_discrete(
    labels = c(
      expression(italic("Temora longicornis")),
      expression(italic("Calanus sp.")), 
      expression(italic("Centropages hamatus")),
      expression("Unknown"),
      expression(italic("Paracalanus parvus")),
      expression(italic("Pseudocalanus sp.")),
      expression(italic("Centropages typicus")),
      expression(italic("cf. Microsetella sp."))
    )
  )+
  
  # Add a bar plot layer using actual values as heights, with specified colors
  geom_bar(
    stat = "identity",  # Use actual values for heights
    fill = "grey60",  # Set fill color of bars
    colour = "grey40"  # Set border color of bars
  )+  

  # Add labels for the title, x-axis, and y-axis
  labs(
    title = "Marine, n = 56",  # Set plot title
    x = "",  # Set x-axis label to empty
    y = "Proportion (%)"  # Set y-axis label
  )+
  
  # Add text labels to the bar plot
  geom_text(
    aes(label = Count),  # Use 'Count' for labels
    vjust = -0.4,  # Adjust vertical position of labels
    size = 4,  # Set text size
    color = "black"  # Set text color
  ) +  
  
  # Apply a classic theme to the plot
  theme_classic()+  

  # Customize various aspects of the plot's appearance
  theme(
    axis.text = element_text(size = 9, colour = "black"),  # Customize axis text
    axis.text.x = element_text(size = 9, angle = 50, hjust = 1, face = "italic"),  # Customize x-axis text
    axis.title.y = element_text(size = 12, vjust = 1.8, colour = "black")  # Customize y-axis title
  )
```

```
# Save the plot as an image (optional)
## ggsave("Taxa_comp_Mar.png", width = 100, height = 100, units = "mm", dpi=700)
```

### R session information

```
# Display the session information and format it using the 'pander' package
sessionInfo() %>% 
  pander::pander()
```

**R version 4.2.3 (2023-03-15 ucrt)**

**Platform:** x86\_64-w64-mingw32/x64 (64-bit)

**locale:** *LC\_COLLATE=Swedish\_Sweden.utf8*,
*LC\_CTYPE=Swedish\_Sweden.utf8*,
*LC\_MONETARY=Swedish\_Sweden.utf8*, *LC\_NUMERIC=C* and
*LC\_TIME=Swedish\_Sweden.utf8*

**attached base packages:** *tcltk*,
*stats*, *graphics*, *grDevices*, *utils*,
*datasets*, *methods* and *base*

**other attached packages:** *webshot2(v.0.1.1)*,
*gtsummary(v.1.7.1)*, *chunkhooks(v.0.0.1)*,
*lubridate(v.1.9.2)*, *forcats(v.1.0.0)*,
*stringr(v.1.5.0)*, *dplyr(v.1.1.2)*,
*purrr(v.1.0.1)*, *tidyr(v.1.3.0)*,
*tibble(v.3.2.1)*, *tidyverse(v.2.0.0)*,
*BiodiversityR(v.2.15-1)*, *knitr(v.1.42)*,
*ggh4x(v.0.2.4)*, *rstatix(v.0.7.2)*,
*ggpubr(v.0.6.0)*, *ggplot2(v.3.4.2)*,
*readr(v.2.1.4)*, *vegan(v.2.6-4)*,
*lattice(v.0.20-45)*, *permute(v.0.9-7)*,
*devtools(v.2.4.5)*, *usethis(v.2.1.6)*,
*MicEco(v.0.9.19)* and *pacman(v.0.5.1)*

**loaded via a namespace (and not attached):**
*utf8(v.1.2.3)*, *tidyselect(v.1.2.0)*,
*lme4(v.1.1-33)*, *htmlwidgets(v.1.6.2)*,
*grid(v.4.2.3)*, *munsell(v.0.5.0)*,
*codetools(v.0.2-19)*, *miniUI(v.0.1.1.1)*,
*withr(v.2.5.0)*, *colorspace(v.2.1-0)*,
*Biobase(v.2.58.0)*, *phyloseq(v.1.42.0)*,
*highr(v.0.10)*, *rstudioapi(v.0.14)*,
*stats4(v.4.2.3)*, *ggsignif(v.0.6.4)*,
*labeling(v.0.4.2)*, *bbmle(v.1.0.25)*,
*GenomeInfoDbData(v.1.2.9)*, *bit64(v.4.0.5)*,
*farver(v.2.1.1)*, *pheatmap(v.1.0.12)*,
*rhdf5(v.2.42.0)*, *vctrs(v.0.6.2)*,
*generics(v.0.1.3)*, *xfun(v.0.39)*,
*timechange(v.0.2.0)*, *markdown(v.1.6)*,
*R6(v.2.5.1)*, *GenomeInfoDb(v.1.34.9)*,
*bitops(v.1.0-7)*, *rhdf5filters(v.1.10.0)*,
*cachem(v.1.0.8)*, *vroom(v.1.6.3)*,
*promises(v.1.2.0.1)*, *scales(v.1.2.1)*,
*nnet(v.7.3-18)*, *gtable(v.0.3.3)*,
*processx(v.3.8.1)*, *sandwich(v.3.0-2)*,
*rlang(v.1.1.1)*, *eulerr(v.7.0.0)*,
*splines(v.4.2.3)*, *broom(v.1.0.4)*,
*checkmate(v.2.2.0)*, *yaml(v.2.3.7)*,
*reshape2(v.1.4.4)*, *abind(v.1.4-5)*,
*backports(v.1.4.1)*, *httpuv(v.1.6.10)*,
*Hmisc(v.5.1-0)*, *tools(v.4.2.3)*,
*Rcmdr(v.2.8-0)*, *ellipsis(v.0.3.2)*,
*jquerylib(v.0.1.4)*, *biomformat(v.1.26.0)*,
*RColorBrewer(v.1.1-3)*, *proxy(v.0.4-27)*,
*BiocGenerics(v.0.44.0)*, *sessioninfo(v.1.2.2)*,
*Rcpp(v.1.0.10)*, *plyr(v.1.8.8)*,
*base64enc(v.0.1-3)*, *zlibbioc(v.1.44.0)*,
*RCurl(v.1.98-1.12)*, *ps(v.1.7.5)*,
*prettyunits(v.1.1.1)*, *rpart(v.4.1.19)*,
*urlchecker(v.1.0.1)*, *S4Vectors(v.0.36.2)*,
*zoo(v.1.8-12)*, *haven(v.2.5.2)*,
*cluster(v.2.1.4)*, *fs(v.1.6.2)*,
*survey(v.4.2-1)*, *magrittr(v.2.0.3)*,
*data.table(v.1.14.8)*, *effects(v.4.2-2)*,
*mvtnorm(v.1.1-3)*, *pkgload(v.1.3.2)*,
*hms(v.1.1.3)*, *RcmdrMisc(v.2.7-2)*,
*mime(v.0.12)*, *evaluate(v.0.21)*,
*xtable(v.1.8-4)*, *readxl(v.1.4.2)*,
*IRanges(v.2.32.0)*, *gridExtra(v.2.3)*,
*compiler(v.4.2.3)*, *bdsmatrix(v.1.3-6)*,
*gt(v.0.9.0)*, *crayon(v.1.5.2)*,
*websocket(v.1.4.1)*, *minqa(v.1.2.5)*,
*htmltools(v.0.5.5)*, *mgcv(v.1.8-42)*,
*later(v.1.3.1)*, *tzdb(v.0.3.0)*,
*Formula(v.1.2-5)*, *snow(v.0.4-4)*,
*DBI(v.1.1.3)*, *MASS(v.7.3-60)*,
*broom.helpers(v.1.13.0)*, *boot(v.1.3-28.1)*,
*relimp(v.1.0-5)*, *Matrix(v.1.5-4)*,
*ade4(v.1.7-22)*, *car(v.3.1-2)*, *cli(v.3.6.1)*,
*mitools(v.2.4)*, *parallel(v.4.2.3)*,
*insight(v.0.19.1)*, *igraph(v.1.4.2)*,
*pkgconfig(v.2.0.3)*, *numDeriv(v.2016.8-1.1)*,
*foreign(v.0.8-84)*, *xml2(v.1.3.4)*,
*foreach(v.1.5.2)*, *bslib(v.0.4.2)*,
*picante(v.1.8.2)*, *chromote(v.0.1.2)*,
*multtest(v.2.54.0)*, *XVector(v.0.38.0)*,
*callr(v.3.7.3)*, *digest(v.0.6.31)*,
*Biostrings(v.2.66.0)*, *rmarkdown(v.2.21)*,
*cellranger(v.1.1.0)*, *htmlTable(v.2.4.1)*,
*nortest(v.1.0-4)*, *commonmark(v.1.9.0)*,
*shiny(v.1.7.4)*, *nloptr(v.2.0.3)*,
*lifecycle(v.1.0.3)*, *nlme(v.3.1-162)*,
*jsonlite(v.1.8.4)*, *Rhdf5lib(v.1.20.0)*,
*carData(v.3.0-5)*, *fansi(v.1.0.4)*,
*labelled(v.2.11.0)*, *pillar(v.1.9.0)*,
*fastmap(v.1.1.1)*, *pkgbuild(v.1.4.0)*,
*survival(v.3.5-5)*, *glue(v.1.6.2)*,
*remotes(v.2.4.2)*, *iterators(v.1.0.14)*,
*pander(v.0.6.5)*, *bit(v.4.0.5)*,
*tcltk2(v.1.2-11)*, *class(v.7.3-22)*,
*stringi(v.1.7.12)*, *sass(v.0.4.6)*,
*profvis(v.0.3.8)*, *doSNOW(v.1.0.20)*,
*memoise(v.2.0.1)*, *e1071(v.1.7-13)* and
*ape(v.5.7-1)*

### Package references

```
# Display citations for multiple packages
citation("pacman")
```

```
## 
## To cite pacman in publications, please use:
## 
##   Rinker, T. W. & Kurkiewicz, D. (2017). pacman: Package Management for
##   R. version 0.5.0. Buffalo, New York. http://github.com/trinker/pacman
## 
## A BibTeX entry for LaTeX users is
## 
##   @Manual{,
##     title = {{pacman}: {P}ackage Management for {R}},
##     author = {Tyler W. Rinker and Dason Kurkiewicz},
##     address = {Buffalo, New York},
##     note = {version 0.5.0},
##     year = {2018},
##     url = {http://github.com/trinker/pacman},
##   }
```

```
citation("devtools")
```

```
## 
## To cite package 'devtools' in publications use:
## 
##   Wickham H, Hester J, Chang W, Bryan J (2022). _devtools: Tools to
##   Make Developing R Packages Easier_. R package version 2.4.5,
##   <https://CRAN.R-project.org/package=devtools>.
## 
## A BibTeX entry for LaTeX users is
## 
##   @Manual{,
##     title = {devtools: Tools to Make Developing R Packages Easier},
##     author = {Hadley Wickham and Jim Hester and Winston Chang and Jennifer Bryan},
##     year = {2022},
##     note = {R package version 2.4.5},
##     url = {https://CRAN.R-project.org/package=devtools},
##   }
```

```
citation("vegan")
```

```
## 
## To cite package 'vegan' in publications use:
## 
##   Oksanen J, Simpson G, Blanchet F, Kindt R, Legendre P, Minchin P,
##   O'Hara R, Solymos P, Stevens M, Szoecs E, Wagner H, Barbour M,
##   Bedward M, Bolker B, Borcard D, Carvalho G, Chirico M, De Caceres M,
##   Durand S, Evangelista H, FitzJohn R, Friendly M, Furneaux B, Hannigan
##   G, Hill M, Lahti L, McGlinn D, Ouellette M, Ribeiro Cunha E, Smith T,
##   Stier A, Ter Braak C, Weedon J (2022). _vegan: Community Ecology
##   Package_. R package version 2.6-4,
##   <https://CRAN.R-project.org/package=vegan>.
## 
## A BibTeX entry for LaTeX users is
## 
##   @Manual{,
##     title = {vegan: Community Ecology Package},
##     author = {Jari Oksanen and Gavin L. Simpson and F. Guillaume Blanchet and Roeland Kindt and Pierre Legendre and Peter R. Minchin and R.B. O'Hara and Peter Solymos and M. Henry H. Stevens and Eduard Szoecs and Helene Wagner and Matt Barbour and Michael Bedward and Ben Bolker and Daniel Borcard and Gustavo Carvalho and Michael Chirico and Miquel {De Caceres} and Sebastien Durand and Heloisa Beatriz Antoniazi Evangelista and Rich FitzJohn and Michael Friendly and Brendan Furneaux and Geoffrey Hannigan and Mark O. Hill and Leo Lahti and Dan McGlinn and Marie-Helene Ouellette and Eduardo {Ribeiro Cunha} and Tyler Smith and Adrian Stier and Cajo J.F. {Ter Braak} and James Weedon},
##     year = {2022},
##     note = {R package version 2.6-4},
##     url = {https://CRAN.R-project.org/package=vegan},
##   }
```

```
citation("readr")
```

```
## 
## To cite package 'readr' in publications use:
## 
##   Wickham H, Hester J, Bryan J (2023). _readr: Read Rectangular Text
##   Data_. R package version 2.1.4,
##   <https://CRAN.R-project.org/package=readr>.
## 
## A BibTeX entry for LaTeX users is
## 
##   @Manual{,
##     title = {readr: Read Rectangular Text Data},
##     author = {Hadley Wickham and Jim Hester and Jennifer Bryan},
##     year = {2023},
##     note = {R package version 2.1.4},
##     url = {https://CRAN.R-project.org/package=readr},
##   }
```

```
citation("ggpubr")
```

```
## 
## To cite package 'ggpubr' in publications use:
## 
##   Kassambara A (2023). _ggpubr: 'ggplot2' Based Publication Ready
##   Plots_. R package version 0.6.0,
##   <https://CRAN.R-project.org/package=ggpubr>.
## 
## A BibTeX entry for LaTeX users is
## 
##   @Manual{,
##     title = {ggpubr: 'ggplot2' Based Publication Ready Plots},
##     author = {Alboukadel Kassambara},
##     year = {2023},
##     note = {R package version 0.6.0},
##     url = {https://CRAN.R-project.org/package=ggpubr},
##   }
```

```
citation("rstatix")
```

```
## 
## To cite package 'rstatix' in publications use:
## 
##   Kassambara A (2023). _rstatix: Pipe-Friendly Framework for Basic
##   Statistical Tests_. R package version 0.7.2,
##   <https://CRAN.R-project.org/package=rstatix>.
## 
## A BibTeX entry for LaTeX users is
## 
##   @Manual{,
##     title = {rstatix: Pipe-Friendly Framework for Basic Statistical Tests},
##     author = {Alboukadel Kassambara},
##     year = {2023},
##     note = {R package version 0.7.2},
##     url = {https://CRAN.R-project.org/package=rstatix},
##   }
```

```
citation("ggh4x")
```

```
## 
## To cite package 'ggh4x' in publications use:
## 
##   van den Brand T (2023). _ggh4x: Hacks for 'ggplot2'_. R package
##   version 0.2.4, <https://CRAN.R-project.org/package=ggh4x>.
## 
## A BibTeX entry for LaTeX users is
## 
##   @Manual{,
##     title = {ggh4x: Hacks for 'ggplot2'},
##     author = {Teun {van den Brand}},
##     year = {2023},
##     note = {R package version 0.2.4},
##     url = {https://CRAN.R-project.org/package=ggh4x},
##   }
```

```
citation("knitr")
```

```
## 
## To cite the 'knitr' package in publications use:
## 
##   Yihui Xie (2023). knitr: A General-Purpose Package for Dynamic Report
##   Generation in R. R package version 1.42.
## 
##   Yihui Xie (2015) Dynamic Documents with R and knitr. 2nd edition.
##   Chapman and Hall/CRC. ISBN 978-1498716963
## 
##   Yihui Xie (2014) knitr: A Comprehensive Tool for Reproducible
##   Research in R. In Victoria Stodden, Friedrich Leisch and Roger D.
##   Peng, editors, Implementing Reproducible Computational Research.
##   Chapman and Hall/CRC. ISBN 978-1466561595
## 
## To see these entries in BibTeX format, use 'print(<citation>,
## bibtex=TRUE)', 'toBibtex(.)', or set
## 'options(citation.bibtex.max=999)'.
```

```
citation("BiodiversityR")
```

```
## 
## To cite BiodiversityR in publications use:
## 
##   Kindt, R. & Coe, R. (2005) Tree diversity analysis. A manual and
##   software for common statistical methods for ecological and
##   biodiversity studies. World Agroforestry Centre (ICRAF), Nairobi.
##   ISBN 92-9059-179-X.
## 
## A BibTeX entry for LaTeX users is
## 
##   @Book{,
##     title = {Tree diversity analysis. A manual and software for common statistical methods for ecological and biodiversity studies},
##     author = {R. Kindt and R. Coe},
##     publisher = {World Agroforestry Centre (ICRAF)},
##     address = {Nairobi (Kenya)},
##     year = {2005},
##     note = {ISBN 92-9059-179-X},
##     url = {http://www.worldagroforestry.org/output/tree-diversity-analysis},
##   }
```

```
citation("tidyverse")
```

```
## 
## To cite package 'tidyverse' in publications use:
## 
##   Wickham H, Averick M, Bryan J, Chang W, McGowan LD, François R,
##   Grolemund G, Hayes A, Henry L, Hester J, Kuhn M, Pedersen TL, Miller
##   E, Bache SM, Müller K, Ooms J, Robinson D, Seidel DP, Spinu V,
##   Takahashi K, Vaughan D, Wilke C, Woo K, Yutani H (2019). "Welcome to
##   the tidyverse." _Journal of Open Source Software_, *4*(43), 1686.
##   doi:10.21105/joss.01686 <https://doi.org/10.21105/joss.01686>.
## 
## A BibTeX entry for LaTeX users is
## 
##   @Article{,
##     title = {Welcome to the {tidyverse}},
##     author = {Hadley Wickham and Mara Averick and Jennifer Bryan and Winston Chang and Lucy D'Agostino McGowan and Romain François and Garrett Grolemund and Alex Hayes and Lionel Henry and Jim Hester and Max Kuhn and Thomas Lin Pedersen and Evan Miller and Stephan Milton Bache and Kirill Müller and Jeroen Ooms and David Robinson and Dana Paige Seidel and Vitalie Spinu and Kohske Takahashi and Davis Vaughan and Claus Wilke and Kara Woo and Hiroaki Yutani},
##     year = {2019},
##     journal = {Journal of Open Source Software},
##     volume = {4},
##     number = {43},
##     pages = {1686},
##     doi = {10.21105/joss.01686},
##   }
```

```
citation("gtsummary")
```

```
## 
## To cite gtsummary in publications use:
## 
##   Sjoberg DD, Whiting K, Curry M, Lavery JA, Larmarange J. Reproducible
##   summary tables with the gtsummary package. The R Journal
##   2021;13:570–80. https://doi.org/10.32614/RJ-2021-053.
## 
## A BibTeX entry for LaTeX users is
## 
##   @Article{gtsummary,
##     author = {Daniel D. Sjoberg and Karissa Whiting and Michael Curry and Jessica A. Lavery and Joseph Larmarange},
##     title = {Reproducible Summary Tables with the gtsummary Package},
##     journal = {{The R Journal}},
##     year = {2021},
##     url = {https://doi.org/10.32614/RJ-2021-053},
##     doi = {10.32614/RJ-2021-053},
##     volume = {13},
##     issue = {1},
##     pages = {570-580},
##   }
```

```
citation("webshot2")
```

```
## 
## To cite package 'webshot2' in publications use:
## 
##   Chang W (2023). _webshot2: Take Screenshots of Web Pages_. R package
##   version 0.1.1, <https://CRAN.R-project.org/package=webshot2>.
## 
## A BibTeX entry for LaTeX users is
## 
##   @Manual{,
##     title = {webshot2: Take Screenshots of Web Pages},
##     author = {Winston Chang},
##     year = {2023},
##     note = {R package version 0.1.1},
##     url = {https://CRAN.R-project.org/package=webshot2},
##   }
```

```
citation("MicEco")
```

```
## 
## To cite package 'MicEco' in publications use:
## 
##   Russel J (2023). _MicEco: Various functions for microbial community
##   data_. R package version 0.9.19.
## 
## A BibTeX entry for LaTeX users is
## 
##   @Manual{,
##     title = {MicEco: Various functions for microbial community data},
##     author = {Jakob Russel},
##     year = {2023},
##     note = {R package version 0.9.19},
##   }
```
